# Genetics and material culture support repeated expansions into Paleolithic Eurasia from a population hub out of Africa

**DOI:** 10.1101/2021.05.18.444621

**Authors:** Leonardo Vallini, Giulia Marciani, Serena Aneli, Eugenio Bortolini, Stefano Benazzi, Telmo Pievani, Luca Pagani

## Abstract

The population dynamics that followed the out of Africa expansion (OoA) and the whereabouts of the early migrants before the differentiation that ultimately led to the formation of Oceanian, West and East Eurasian macro populations have long been debated. Shedding light on these events may, in turn, provide clues to better understand cultural evolution in Eurasia between 50kya and 35kya.

Here we analyze Eurasian Paleolithic DNA evidence to provide a comprehensive population model and validate it in light of available material culture. Leveraging on our integrated approach we propose the existence of a Eurasian population Hub, where *Homo sapiens* lived between the OoA and the broader colonization of Eurasia, which was characterized by multiple events of expansion and local extinction. A major population wave out of Hub, of which Ust’Ishim, Bacho Kiro and Tianyuan are unadmixed representatives, is broadly associated with Initial Upper Paleolithic lithics and populated West and East Eurasia before or around 45 kya, before getting largely extinct in Europe. In this light, we suggest a parsimonious placement of Oase1 as an individual related to Bacho Kiro who experienced additional Neanderthal introgression. Another expansion, started before 38 kya, is broadly associated with Upper Paleolithic industries and repopulated Europe with sporadic admixtures with the previous wave (GoyetQ116-1) and more systematic ones while moving through Siberia (Yana, Mal’ta).

## Introduction

Several layers of genetic, palaeoanthropology and palaeoclimatology evidence point to 70-60 kya as the most likely time window for the major colonization of Eurasia by *Homo sapiens*. For several millennia, however, the human population Out of Africa (OoA) did not expand much from a demographic perspective, with the divergence between Eastern and Western Eurasian populations inferred not earlier than 45-40 kya ago from modern DNA data(Bergström et al., 2020; Choin et al., 2021; Malaspinas et al., 2016; Pagani et al., 2015, 2016; Schiffels & Durbin, 2014; Soares et al., 2012). We can imagine that, after leaving Africa(Pagani, 2019), the ancestors of all non Africans lived somewhere on the new continent, interbred with Neanderthals(Green et al., 2010) and persisted as a single population for at least 15 thousand years (conservatively, the time between the OoA bottleneck and the split between European and East Asian populations, marking the beginning of a broader expansion) and later expanded from this “population Hub” ultimately colonizing all of Eurasia and further.

Disentangling the processes underlying population and technological shifts between ∼50 kya and ∼35 kya during the broader colonization of Eurasia is critical to explain the shaping of current *H. sapiens* genetic diversity out of Africa, and to understand if cultural change documented in the archaeological record can be attributed to population movements, human interactions, convergence or any intermediate mechanism of biocultural exchange. This time interval is characterised by the appearance and turnover of several techno-complexes which, based on technological characteristics, we divide into: i) production of blades using volumetric and levallois methods here extensively defined as Initial Upper Paleolithic (IUP) (E. Boëda et al., 2013; Kuhn, 2019; Kuhn & Zwyns, 2014) ; ii) lithic industries characterized by the production of blades and bladelets often together with ornaments and bone tools and here inclusively defined as Upper Paleolithic (UP) ; iii) non-Mousterian and non-IUP technologies appeared during the Middle to Upper Palaeolithic transition, comprising Uluzzian, Châtelperronian, Szeletian and Lincombian-Ranisian-Jermanowician (LRJ) (see Supplementary Section 1 for an in-depth definition of these material culture labels). Only a few contexts present with both material culture and stratigraphically related human remains for which aDNA is available (Table 1, Table S1), which leads to many possible scenarios of association between cultural change and human migration, as well as with inter- and intra-specific human interaction.

**Table 1.** Summary of Paleolithic individuals used for qpGraph modeling, see table S1 for full details

| Sample | Techno-complex | Country | Date(kya) | Reference |
| --- | --- | --- | --- | --- |
| Zlatý Kůň | Szeletian/IUP<sup>a</sup> | Czech Republic | >45 | (Prüfer et al., 2021) |
| Ust'Ishim | IUP<sup>a</sup> | Russia | 45 | (Fu et al., 2014) |
| Bacho Kiro | IUP | Bulgaria | 45 | (Hajdinjak et al., 2021) |
| Oase1 | IUP/UP<sup>a</sup> | Romania | 40 | (Fu et al., 2015) |
| Tianyuan | IUP<sup>a</sup> | China | 40 | (Yang et al., 2017) |
| Kostenki14 | UP<sup>a</sup> | Russia | 38 | (Fu et al., 2016) |
| GoyetQ116-1 | UP<sup>a</sup> | Belgium | 35 | (Fu et al., 2016) |
| Sunghir | UP | Russia | 34 | (Sikora et al., 2017) |
| Yana | UP | Russia | 32 | (Sikora et al., 2019) |
| Mal'ta | UP | Russia | 24 | (Raghavan et al., 2014) |

a: attribution based on nearby coeval sites

From a genetic perspective, two recent studies found that around 45 kya, nearby European territories were occupied by either a human population basal to all Eurasians: Zlatý Kůň(Prüfer et al., 2021), or by a human population that is closer to ancient and contemporary East Asians than to later and contemporary Europeans(Hajdinjak et al., 2021), which triggers the question of where and how the European component, later represented by Kostenki14 (38 kya), differentiated after separating from East Asians.

## Results

We used Treemix(Pickrell & Pritchard, 2012) for a preliminary, unsupervised exploration of the genetic relationships among the crucial samples listed in Table 1 and, after adding up to 5 admixture events we found that it fails in identifying well known events such as the Neanderthal admixture shared by all Eurasians that is instead monopolized by Bacho Kiro, and noted high residuals in any solution offered by this method (results not shown). Despite known limitations involved in its reliance on initial topologies based on the operator’s assumptions, we resorted to qpGraph (Supplementary Section **2**), implementing the simple population tree proposed by Pruefer and colleagues(Prüfer et al., 2021) and proceeded step-wise to minimize the aforementioned issues. We tried to add Bacho Kiro in all plausible positions without invoking additional admixture events, with the exception of the extra Neanderthal introgression already documented for the Bacho Kiro samples(Hajdinjak et al., 2021) (Figure S1, Supplementary Section 3). To allow the likely European Neanderthal population that admixed with the ancestors of Bacho Kiro to be different from the Neanderthal population that admixed with the ancestors of all non Africans shortly after the Out of Africa, we modeled it as a different node closer to the Neanderthal from Vindija (Croatia); if it was indeed the same Neanderthal population the inferred drift between the two nodes would be 0. Additionally, to avoid our results to be driven by later population interaction between Eurasia and Western Africa(Chen et al., 2020) or by the putative admixture of Mbuti pygmies with an archaic ghost hominin(Hammer et al., 2011; Hsieh et al., 2016; Lipson et al., 2020) we used four ancient South African hunter-gatherers(Schlebusch et al., 2017; Skoglund et al., 2017) (Table S1) instead of Mbuti, and found the best placement of Bacho Kiro as a sister group of Tianyuan (Figure S1.C, Supplementary Section 3.2). This placement is different from the one proposed by(Hajdinjak et al., 2021) and we found little support for a model placing Bacho Kiro as an early branch among the Out of Africa lineage (Figure S1.A) when immediately accounting for its higher Neanderthal ancestry, possibly also thanks to the availability of Zlatý Kůň, who may provide a good guidance to the basal OoA genetic landscape. After adding Kostenki14 as a key ancient European sample, we found that Ust’Ishim would fit better as a basal split along the branch leading to Tianyuan and Bacho Kiro (Supplementary Section **3.3**) (Figure S2).

The scenario emerging from our proposed tree (Figure 1A, Figure S2.B) depicts Zlatý Kůň as a population basal to all subsequent splits within Eurasia. Downstream of Zlatý Kůň, the separation of Ust’Ishim and other genetically East Asian ancient samples (Bacho Kiro and Tianyuan, red in Figure 1A) from the branch eventually leading to Kostenki and Sunghir (genetically West Eurasians with a deep shared genetic drift - 49 units -, in blue in Figure 1A) defines the first major subdivision of Eurasian genetic components. Although this represents only one of several otherwise unexplored possible trees, its topology is broadly matched by the spatiotemporal distribution of material cultural evidence at a cross-continental scale, according to the present state of archaeological knowledge. From a chronological point of view the right branch (red) of Figure 1A presents samples dated ∼45/40 kya, while the left (blue) one is instead characterised by a younger date of the Kostenki and Sunghir samples (38 and 34 kya respectively). The structure emerging from genetic distances is also supported and confirmed by technological evidence. The earlier, red branch is consistently populated by contexts either directly showing or surrounded by geographic and temporal proxies exhibiting IUP technology (Table S1, Table S2). The later, blue branch is instead predominantly characterised by contexts with UP technology (Table S1, Table S2). Finally, the basal Zlatý Kůň is coeval to Eastern European sites exhibiting Mousterian, non-Mousterian and IUP technologies (Table S1, Table S3).

**Figure 1.**
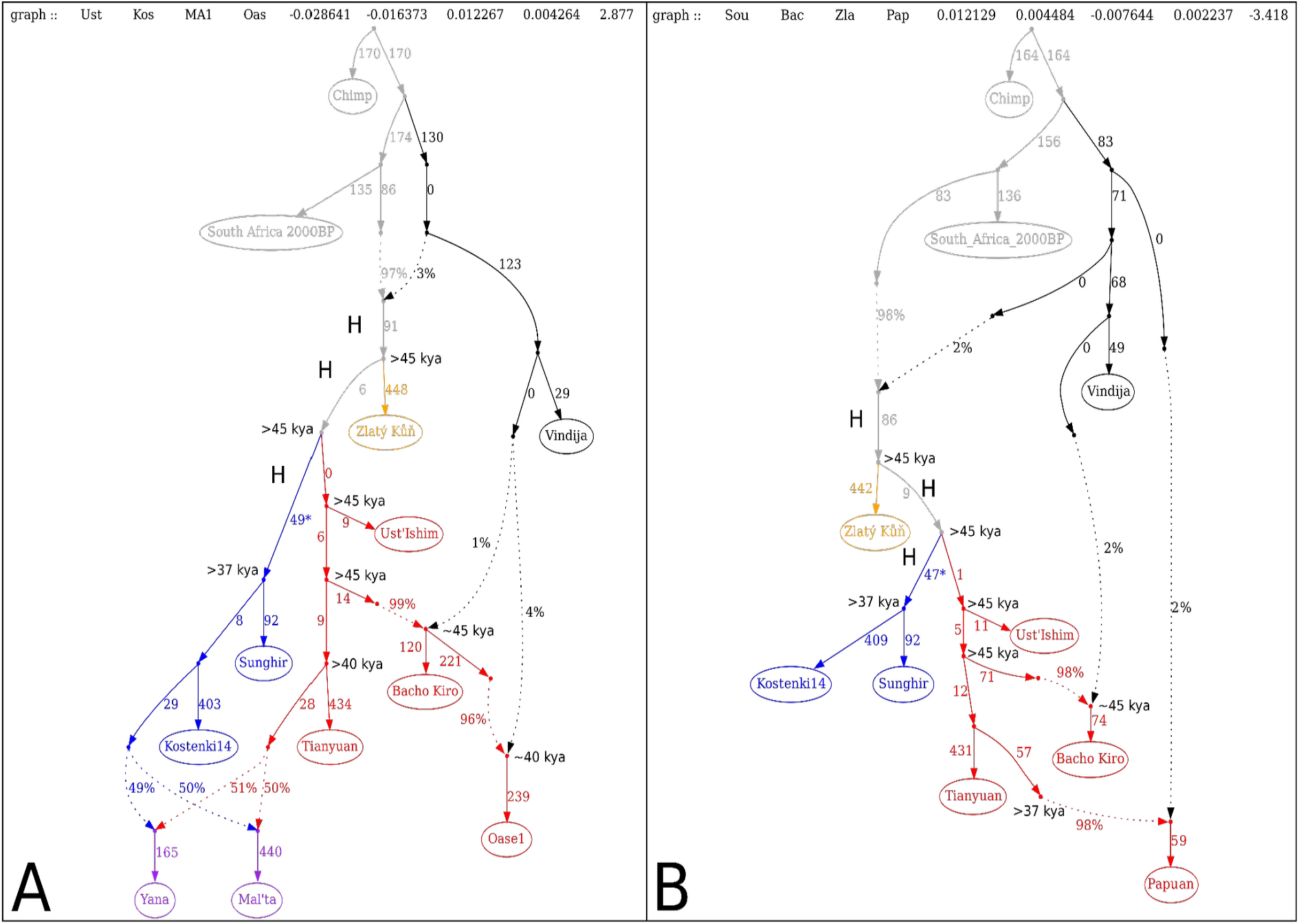
qpGraph models for Paleolithic Eurasia. **Panel A:** Best fit population model that recapitulates the major population streams from an Out of Africa hub, colored according to the most parsimonious lithic culture affiliation (Transitional complexes: yellow; Initial Upper Paleolithic, IUP: red; Upper Paleolithic, UP: blue). See Supplementary Sections 1 and 3 for more details on the qpGraph generation and on the material culture labels. The tree proposed here is based on 71539 SNPs due to the presence of Oase1, but its significance holds with a greater number of SNPs (302476) when Oase1 is removed (Figure S5.C). **Panel B:** Modern Papuans can be modelled as a terminal branch of the Paleolithic expansion that was associated with IUP in Eurasia. Such a model, based on 417400 SNPs, is just one of the six acceptable possibilities we identified (Figure S7) and is reported here just on the basis of its parsimonious nature. Nodes labelled with “H” represent population differentiation inferred to have happened inside the population hub out of Africa. Asterisks indicate genetic drift putatively occurred inside the hub, which differentiates the West and East Eurasian genetic components.

Notably, Oase1, which has so far been regarded as a lineage unrelated to extant Eurasians(Fu et al., 2015), can be modeled as an additional admixture between the Bacho Kiro population, or a closely related one, and Neanderthals (Figure 1A, Supplementary Section 3.4). This result is coherent from a geographical and chronological perspective, being Oase1 dated to around 5 ky later with respect to Bacho Kiro and being located just a few hundred kilometers away. Incidentally, the proposed placement of Oase1 on the graph provides support for the claim made in the original Oase1 publication(Fu et al., 2015) about an additional pulse of Neanderthal admixture experienced by the Oase1 ancestors between the one shared by all non Africans and the one that occurred 4-6 generations before it lived. We here propose that such an event may be shared with the Bacho Kiro population or with a closely related one, which could be seen as ancestral to Oase1. To rule out the possibility that Oase1 attraction to Bacho Kiro is driven by the excess of Neanderthal ancestry they share, we masked the most recent Neanderthal introgressed segments of Oase1 and re-run the analysis: Oase1 holds its position and can be modeled without the last admixture event with Neandethal hence confirming its genuine affinity with Bacho Kiro (Figure S3.D, Supplementary section 3.4). Finally, this placement for Oase1 is consistent with the reported East Asian genetic affinities for another sample from the same site (Oase2, Supplementary Section 3.4).

Our model inferred by genetic data and supported by material culture, would then explain Ust’Ishim as the result of early IUP movements towards Siberia(Nicolas Zwyns, 2021; Nicolas Zwyns et al., 2019), and the presence of Bacho Kiro-like populations in Europe at least from 45kya as part of a broader peopling event that reached as far East as Tianyuan (40kya) with little or no interaction with pre-existing Zlatý Kůň-like groups (Figure S4, Supplementary Section 3.5) but with occasional contacts with Neanderthals(Fu et al., 2015; Hajdinjak et al., 2021). The UP branch in the model would have then emerged from a putative OoA population Hub well after 45kya, a scenario that finds support in previous hypotheses on the appearance of UP techno-complexes (e.g. Aurignacian) in Europe, although the role of migrations and exchange between Europe, and Western Asia and the Levant is still debated(Bataille et al., 2020; Conard, 2002; Davies, 2007; Falcucci et al., 2020; Hoffecker, 2009; Hublin, 2015; Le Brun-Ricalens et al., 2009; Mellars, 2006; Nigst et al., 2014; Teyssandier, 2006; Zilhão, 2014). The origin and spatiotemporal development of later UP techno-complexes (e.g. Gravettian) are also still debated, and are currently interpreted as the outcome of mixed processes involving regional adaptation, functional convergence, and exchange between populations which took place in Europe(Kozłowski, 2015; Moreau, 2012; M. Otte, 2013; Marcel Otte, 2017; Pesesse, 2010) after the putative split from the OoA hub. As far as Northern Asia is concerned, the UP legacy may be responsible for the West Eurasian components already reported in ancient Siberian samples dated between 34kya (Salkhit(Massilani et al., 2020)), 31kya (Yana1(Sikora et al., 2019)) and 24kya (Mal’ta(Raghavan et al., 2014)). Admixture events in varying proportions between sister groups of Sunghir and Tianyuan can indeed explain this observation (Figure 1A, purple leaves; Figure S5, Supplementary Section 3.6). This is further supported by the younger chronology for these two sites which is compatible with a stepwise arrival of West Eurasian components in Siberia following the UP exit from the Hub sometimes before 38 kya. The lack of West Eurasian components in Tianyuan and in subsequent East Asian individuals may provide clues on the resistance of those groups to the incoming UP population movements, or on subsequent re-expansion from a genetically IUP-like population reservoir.

West Eurasian IUP populations, on the other hand, likely declined and ultimately disappeared, as suggested by the fact that our population tree is compatible with the arrival in Europe of UP groups who experienced no further admixture with pre-existing IUP or Neanderthals, with the exception of GoyetQ116-1. The East Asian component carried by this sample(Fu et al., 2016) can be described as an interaction between pre-existing Bacho Kiro-like and incoming UP groups in West Europe with an additional Tianyuan-like component. Such a variegated East Asian substrate found in the otherwise West Eurasian GoyetQ116-1 sample accounts for yet undescribed complexities within the IUP population branch (Figure S6, Supplementary Section 3.7). Interestingly, the fade of the IUP populations in Europe coincides with the extinction of the last Neanderthals(Hajdinjak et al., 2018).

Given the relatively simple population tree needed to explain the post OoA Eurasian population movements using aDNA samples available to date, and benefiting from the basal position of Zlatý Kůň, we tried to model Oceanian populations (using modern Papuans) within the emerging picture to resolve a long lasting debate on their topological position with respect to East and West Eurasians. Starting from the topologies proposed in(Choin et al., 2021; Malaspinas et al., 2016) we first tried to place Papuans as the most basal branch along the non-African sub-tree, allowing for the documented Denisova admixture(Meyer et al., 2012; Reich et al., 2011). We avoided including the sampled Denisova aDNA within the population tree to eliminate attractions from yet uncharacterized, deep splits along the hominin branch(Hajdinjak et al., 2021), and opted for letting qpGraph infer Denisova as a basal split along the archaic human lineage. Simply placing Papuans in a basal position (either before or after the split of Zlatý Kůň) was rejected and highlighted a notable attraction between them and Tianyuan. We then modelled Papuans as sister of Tianyuan (Figure 1.B, Figure S7.A)(Wall, 2017), a solution that yielded only two marginal outliers that could be resolved when taking all the SNPs into account (Supplementary Section 3.8). This topology could be further improved when allowing for a contribution of a basal lineage, whose magnitude decreased the deeper was its position along and beyond the OoA tree: 94% as the ancestor of Bacho Kiro and Tianyuan (Figure S7.B) or 53% as the most basal IUP lineage (Figure S7.C), 40% before the West/East Eurasia split (Figure S7.D), 2% before the Zlatý Kůň lineage, along the OoA path (Figure S7.E) and 2% as an extinct OoA (xOoA, Figure S7.F) as proposed by Pagani and colleagues(Pagani et al., 2016). Notably, all the acceptable solutions for the placement of Papuans within the broader OoA tree encompass a Denisova contribution and confirm Zlatý Kůň as the most basal human genome among the ones ever found Out of Africa. Taken together with a lower bound of the final settlement of Sahul at 37 kya (the date of the deepest population splits estimated by(Malaspinas et al., 2016)) it is reasonable to describe Papuans as either an almost even mixture between East Asians and a lineage basal to West and East Asians occurred sometimes between 45 and 38kya, or as a sister lineage of East Asians with or without a minor basal OoA or xOoA contribution. We here chose to parsimoniously describe Papuans as a simple sister group of Tianyuan, cautioning that this may be just one out of six equifinal possibilities.

## Discussion

In conclusion, via introducing the concept of post OoA population Hub, our model provides a robust, elegant and parsimonious framework to explain the relationships between the most representative ancient human DNA available to date between 60 and 24kya across Eurasia, which can be accommodated within just three expansions out of a putative OoA Hub (Figure 2, Supplementary Section 4). Zlatý Kůň may represent an early expansion, which left little to no traces in subsequent Eurasians and occurred before or around 45kya (Figure 2A). We speculate this population movement might be linked either to IUP(Richter et al., 2008; Tostevin, 2003) or to non-Mousterian and non-IUP lithic techno-complexes that appeared in Central and Eastern Europe between 48-44kya (e.g., Szeletian, LRJ)(Nejman et al., 2017; Svoboda, 2000).

**Figure 2.**
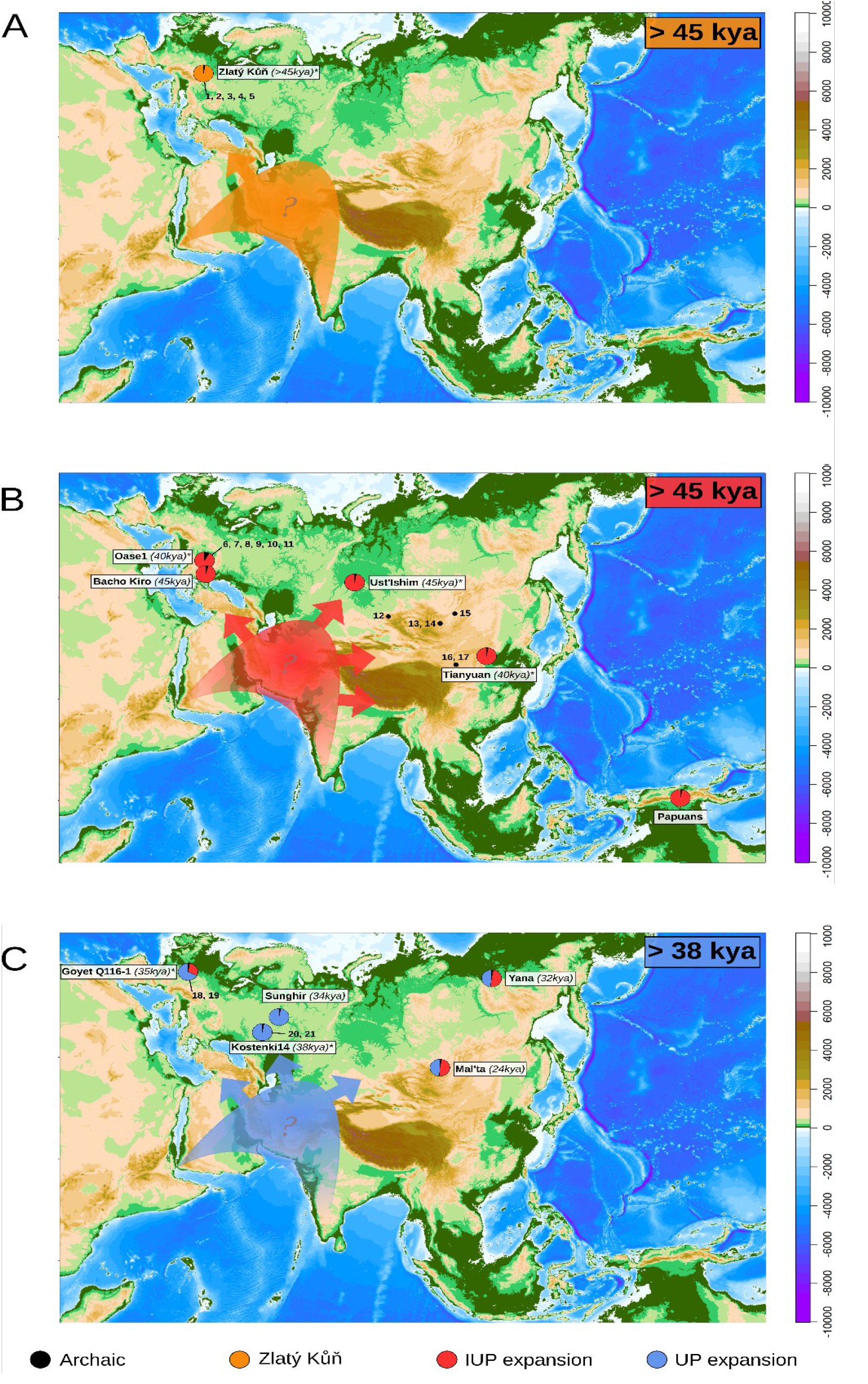
Subsequent expansions into Eurasia from a population hub out of Africa. **Panel A**: Zlatý Kůň can be described as a putative early expansion from the population formed after the major expansion Out of Africa and hybridization with Neanderthals, and could be linked with transitional cultures found in Europe 48-45 kya. **Panel B:** Representative samples dated between 45 kya and 40 kya across Eurasia can be ascribed to a population movement with uniform genetic features and material culture consistent with an Initial Upper Palaeolithic affiliation and which can also explain Oase1 after allowing for additional Neanderthal contributions; modern Papuans may be genetically seen as an extreme extension of this movement. **Panel C:** following local genetic differentiation, a subsequent population expansion could explain the genetic components found in ancient samples <38 kya which contain it in unadmixed form (Kostenki14, Sunghir) or admixed with pre-existing IUP components (Goyet Q116-1, Yana1, Mal’ta). The dates at the top right of each map provide a lower bound, based on the C14 of the earliest available sample for the inferred population wave. * indicate sites for which material culture was not available in direct association. For these sites the nearest spatio-temporal proxies were used, as indicated in Table S1. Numbers on the map refer to the position of relevant proxies: 1: Szeleta (S); 2: Pod_Hradem (S); 3: Moravský_Krumlov_IV (S); 4: Stranska_Skala_III-IIIc (IUP); 5: Brno-Bohunice (IUP); 6: Bacho_Kiro_IUP_layer_11 (IUP); 7: Ořechov_IV_–_Kabáty (IUP); 8: Brno-Bohunice (IUP); 9: Românesti-Dumbravita (UP); 10: Cosava (UP); 11: Tincova (UP); 12: Kara_Bom_OH_5_OH6 (IUP); 13: Tolbor-4_layer_4-5-6 (IUP); 14: Tolbor-16_layer_6 (IUP); 15: Kamenka_A (IUP); 16: Suindonggou_1 (IUP); 17: Suindonggou_2 (IUP); 18: Maisières-Canal (UP); 19: Spy_Ossiferous_Horizon_2 (UP); 20: Kostenki_12_Vokov (UP); 21: Kostenki_1 (UP).

A subsequent expansion (linked to IUP in Eurasia) can be dated earlier than 45kya as proposed by Zwyn and colleagues(Nicolas Zwyns et al., 2019), and here we propose it to be a wider phenomenon that populated the broad geographic area between Mediterranean Levant(Eric Boëda & Bonilauri, 2006; Kadowaki et al., 2021; Kuhn et al., 2009; Leder, 2017; Marks & Kaufman, 1983), East Europe(Fewlass et al., 2020; Hublin et al., 2020; Richter et al., 2008), Siberia-Mongolia(Derevianko et al., 2013; Kuhn, 2019; Rybin et al., 2020; N. Zwyns et al., 2012; Nicolas Zwyns et al., 2019; N. Zwyns & Lbova, 2019) and East Asia(E. Boëda et al., 2013; Morgan et al., 2014; Peng et al., 2020) in less than 5ky, reaching as far South as Papua New Guinea before 38kya, and which eventually died out in Europe after repeated admixtures with Neanderthals (Bacho Kiro and Oase1 being two notable examples) (Figure 2.B). In Western Europe, in the same timeframe, this interaction has been suggested as a trigger for the development of Chatelperronian material culture(Roussel et al., 2016), while the Uluzzian techno-complex in Mediterranean Europe may tentatively be better explained by an additional, yet uncharacterized expansion from the Hub(Benazzi et al., 2011; Marciani et al., 2020) although genomic data from Uluzzian strata is still lacking. The Uluzzian technocomplex is indeed characterized by unprecedented versatility and efficient management of production costs, considerably lower standardisation in design compared to Mousterian industries, lower time and energy expenditure for initialisation and management of volumes, and a much shorter response time to change in raw material or environmental conditions(Arrighi, Marciani, et al., 2020; Collina et al., 2020; Marciani et al., 2020; Moroni et al., 2013, 2018; Peresani et al., 2019; Riel-Salvatore, 2007, 2009, 2010; Silvestrini et al., 2021). In addition, this technological change is linked to the appearance of complementary tools(Haidle et al., 2015)^,(Sano et al., 2019)^, innovations in hunting strategies(Boscato & Crezzini, 2012; Romandini et al., 2020), and a shared package of symbolic artefacts(Arrighi, Bortolini, et al., 2020; Arrighi, Moroni, et al., 2020). All these almost unique and distinctive elements make it very difficult to infer heritable continuity with local Mousterian material culture, or preferential technological proximity with any other IUP/UP Eurasian technology, and support the hypothesis of an independent population expansion.

The last major expansion needed to explain the observed data (UP) took place later than 45 kya and before 38kya and repopulated (Kostenki, Sunghir), or interacted with, pre-existing human groups (GoyetQ116-1, Figure S6) in Europe, and admixed with members of the previous IUP wave in Siberia (Yana, Mal’ta and perhaps Salkhit) as it moved East in the subsequent 5-10kya (Figure 2.C). The split time between East and West Eurasians estimated at ∼40 kya(Malaspinas et al., 2016) from modern genomes and the differentiation of these two macro-populations can therefore be explained by the inferred timing of the IUP exit from the Hub, followed by subsequent diversification within the Hub of the ancestors of West Eurasians, later mitigated by ongoing cross-Eurasian gene flow. Importantly, the qpGraph results that form the basis for this emerging picture describe only one of the many possible arrangements of the genetic data we explored and, by itself, it shouldn’t be interpreted as the only or best outcome. Remarkably though, the arrangement described here matches and simplifies scenarios inferred from material culture evidence, and provides a framework for placing genetic and cultural data onto a coherent landscape.

In this paper, we used extensive cultural categories that are in agreement with the demic movements at the present, coarse scale of analysis. The general trends observed here emerge from a broad perspective, and further work is needed to test specific hypotheses concerning actual processes of branching, local adaptation, cultural transmission and convergence(Groucutt, 2020; Kuhn, 2019). Similarly, we remain oblivious about the precise location of the inferred population Hub, although North Africa or West Asia seem the most plausible candidates. More ancient genomes are needed, as well as a better understanding of the role of South and SouthEast Asia, for which currently known material culture suggests complex trajectories(Allen & O’Connell, 2014; Bird et al., 2019; Bradshaw et al., 2021; Clarkson et al., 2017; Dennell & Petraglia, 2012; Michel et al., 2016; O’Connell et al., 2018; Petraglia et al., 2010; Shackelford et al., 2018; Sun et al., 2021; Westaway et al., 2017).

## Material and Methods

We downloaded the aligned sequences for the IUP Bacho Kiro individuals from the European Nucleotide Archive (accession number PRJEB39134), and merged files from the same individual. We clipped 3 bases off the ends of each read to avoid excess of aDNA damage using trimBam (https://genome.sph.umich.edu/wiki/BamUtil:_trimBam) and then generated pseudo-haploid calls with the genotype caller pileupCaller (https://github.com/stschiff/sequenceTools/tree/master/src-pileupCaller) using the majorityCall option that, for each locus, either chooses the allele supported by the highest number of reads or picks one randomly if the number is the same. Only positions with base quality and read quality higher than 30 were considered. The processed Eigenstrat files of the Zlatý Kůň individual were kindly provided by Prof. Cosimo Posth and Dr. He Yu.

We merged the files with the 1240K “Allen Ancient DNA Resource” v44.3 database (https://reich.hms.harvard.edu/allen-ancient-dna-resource-aadr-downloadable-genotypes-present-day-and-ancient-dna-data)(Bergström et al., 2020; Skoglund et al., 2015), then kept only autosomal SNPs (1,150,639) using Plink 1.9(Chang et al., 2015) and then converted back to Eigenstrat format using Admixtools convertf(Patterson et al., 2012).

We ran Admixtools qpGraph(Patterson et al., 2012), https://github.com/DReichLab/AdmixTools, qpGraph version: 7365) with the following parameters: outpop: NULL, blgsize: 0.05, lsqmode: YES, diag: .0001, useallsnps: NO, bigiter: 6, forcezmode: YES, initmix: 10000, precision: .0001, zthresh: 3.0, terse: NO, hires: YES.

## Supporting information

ValliniEtAl_Supplementary_Information

## Data and Code Availability

No new data or software was generated for this study. All software, lithic information and genomes are publicly available from the published sources.

Allen ancient DNA resource data repository: https://reich.hms.harvard.edu/allen-ancient-dna-resource-aadr-downloadable-genotypes-present-day-and-ancient-dna-data

European Nucleotide Archive (accession numbers PRJEB39134; PRJEB39040) Palaeoenvironmental data: https://www.ncei.noaa.gov/

Software: https://github.com/DReichLab/AdmixTools, qpGraph version: 7365

## Acknowledgements

The authors would like to thank Prof. Cosimo Post and Dr. He Yu for having contributed the processed Eigenstrat files of the Zlatý Kůň genome. Site coordinates were obtained from the ROCEEH Out of Africa Database (ROAD) (http://www.roceeh.org), and the work of the data contributors and ROAD community is gratefully acknowledged. This work was supported by: UniPd PRID 2019 (to S.A.) and by the European Research Council (ERC) under the European Union’s Horizon 2020 research and innovation program (grant agreement No 724046 – SUCCESS; http://www.erc-success.eu/ to S.B., G.M., E.B.).

## Authors Contribution

Conceptualization: L.V, L.P; Data Analyses: L.V., G.M, E.B, S.A.; Drafting of the manuscript: L.V., G.M, E.B, L.P.; Interpretation of Results: T.P, S.B., L.P.; Revision of the manuscript: all authors.

