## Supplementary material for "Genetics and material culture support repeated expansions into Paleolithic Eurasia from a population hub out of Africa": ValliniEtAl_Supplementary_Information

Supplementary Materials include:

- Supplementary Information
Section 1: Material Culture — Page 2
Section 2: Genetic Dataset — Page 11
Section 3: qpGraph analyses — Page 14
Section 4: Paleomaps plotting — Page 25

- Supplementary Figures S1 to S7
- Supplementary Tables S1 to S4

### Supplementary Section 1: Material Culture

Many of the DNA samples discussed in this paper come from sites without stratigraphic association with diagnostic archaeological materials. To overcome this issue, we use layers of sites located in proximity of and coeval to DNA sampling sites (Table S1).

**Table S1. Cultural proxies for the ancient genomes analysed in the current work**. M: Mousterian; S: Szeletian; IUP: Initial Upper Paleolithic; UP: Upper Paleolithic; [a] indicates an attribution based on nearby coeval sites, which are specified in the subsequent rows. The coordinates of the sites are taken from the reference articles or from the ROCEEH Out of Africa Database (ROAD) (http://www.roceeh.org).

| DNASample/ Proxy | Sample/Site name | Techno-complex | Country | Coordinates | Date (cal kyr BP) | Ref |
|---|---|---|---|---|---|---|
| **DNASample** | **Vindija** | **M** | **Croatia** | **46.29 N, 16.07 E** | ~50 | * |
| Pre-expansion sites in the Levant | Ksar_Akil_XXII-XXV | IUP | Lebanon | 33.91 N, 35.64 E | ~45-43 | (Bosch et al., 2015; Leder, 2014, 2017) |
| Pre-expansion sites in the Levant | Üçagizli_F,Fa,Fb-c,G,H,H1-3,I | IUP | Turkey | 35.98 N, 35.96 E | ~45-40 | (Kuhn, 2004; Kuhn et al., 2009) |
| Pre-expansion sites in the Levant | Umel'Tlel | IUP | Syria | 35.27 N, 38.89 E | ~40 | (Eric Boëda & Bonilauri, 2006) |
| Pre-expansion sites in the Levant | Boker_Tachtit_layer_4 | IUP | Israel | 30.84 N, 34.78 E | ~45-40 | (Kadowaki et al., 2015; Marks & Kaufman, 1983) |
| **DNASample** | **UstIshim** | **IUP[a]** | **Russia** | **57.7 N, 71.1 E** | ~44 | * |
| Ust_Ishim | Kara_Bom_OH_5_OH6 | IUP | Russia | 50.72 N, 85.57 E | >45 | (N. Zwyns et al., 2012) |
| Ust_Ishim | Tolbor-4_layer_4-5-6 | IUP | Mongolia | 49.29 N, 102.97 E | ~45-35 | (Derevianko et al., 2013; Nicolas Zwyns et al., |


2

| | | | | | | |
|---|---|---|---|---|---|---|
| | | | | | | 2019) |
| Ust_Ishim | Tolbor-16_layer_6 | IUP | Mongolia | 49.23 N, 102.92 E | ~45-40 | (Nicolas Zwyns et al., 2019) |
| Ust_Ishim | Kamenka_A | IUP | Russia | 51.44 N, 108.17 E | ~45-40 | (N. Zwyns & Lbova, 2019) |
| **DNASample** | **Tianyuan** | **IUP**[a] | **China** | **39.66 N, 115.87 E** | ~40 | * |
| Tianyuan | Suindonggou_1 | IUP | China | 38.38 N, 106.51 E | <41 | (E. Boëda et al., 2013; Morgan et al., 2014) |
| Tianyuan | Suindonggou_2 | IUP | China | 38.38 N, 106.51 E | <41 | (Li et al., 2019; Peng et al., 2020) |
| **DNASample** | **Kostenki14** | **UP**[a] | **Russia** | **51.23 N, 39.3 E** | ~38 | * |
| Kostenki14 | Kostenki_12_Vokov | UP | Russia | 51.39 N, 39.05 E | <40 | (A. A. Sinitsyn & Hoffecker, 2006) |
| Kostenki14 | Kostenki_1 | UP | Russia | 51.39 N, 39.05 E | <40 | (Hoffecker et al., 2016) |
| **DNASample** | **Oase1** | **IUP-UP**[a] | **Romania** | **45.12 N, 21.9 E** | ~38 | * |
| Oase1 | Bacho_Kiro_IUP_layer_11 | IUP | Bulgaria | 42.95 N, 25.43 E | >45 | (Hublin et al., 2020; Tsanova, 2008) |
| Oase1 | Ořechov_IV_–_Kabáty | IUP | Czech Republic | ~49.06 N, 16.31 E | ~41-35 (under estim.) | (Yuri E. Demidenko et al., 2020) |

| | | | | | | |
|---|---|---|---|---|---|---|
| Oase1 | Brno-Bohunice | IUP | Czech Republic | ~49.12 N 16.37 E | >45 | (Richter et al., 2008) |
| Oase1 | Românesti-Dumbravita | UP | Romania | 45.49 N, 22.19 E | ~45-40 | (Anghelinu & Niţă, 2014; Sitlivy et al., 2012) |
| Oase1 | Cosava | UP | Romania | 45.51 N, 22.19 E | ~45-40 | (Anghelinu & Niţă, 2014; Sitlivy et al., 2014) |
| Oase1 | Tincova | UP | Romania | 45.33 N, 22.9 E | ~45-40 | (Anghelinu & Niţă, 2014; Sitlivy et al., 2014) |
| **DNASample** | **GoyetQ116-1** | **UP**[a] | **Belgium** | **50.45 N, 5.01 E** | ~35 | * |
| GoyetQ116-1 | Maisières-Canal | UP | Belgium | 50.48 N, 3.98 E | <40 | (Pirson et al., 2012) |
| GoyetQ116-1 | Spy_Ossiferous_Horizon_2 | UP | Belgium | 50.48 N, 4.67 E | <40 | (Pirson et al., 2012) |
| **DNASample** | **Sunghir** | **UP** | **Russia** | **56.18 N, 40.50 E** | ~34 | (Dobrovolskaya et al., 2012; Trinkaus & Buzhilova, 2018) |
| **DNASample** | **Yana** | **UP** | **Russia** | **70.72 N, 135.42 E** | ~31 | (Pitulko et al., 2017) |
| **DNASample** | **Mal'ta (MA1)** | **UP** | **Russia** | **52.9 N, 103.5 E** | ~24 | (Khenzykhenova et al., 2019; Lbova, 2019) |
| **DNASample** | **Bacho_Kiro_IUP_layer_11** | **IUP** | **Bulgaria** | **42.95 N, 25.43 E** | ~45 | (Hublin et al., 2020; Tsanova, |

| | | | | | | |
|---|---|---|---|---|---|---|
| | | | | | | 2008) |
| **DNASample** | **Zlatý_Kůň** | **S-IUP[a]** | **Czech Republic** | **49.37 N, 16.72 E** | >45 | * |
| Zlatý Kůň | Szeleta | S | Hungary | 48.06 N, 20. 37 E | ~45-40 | (Adams, 2009; Hauck et al., 2016; Zs Mester, 2010) |
| Zlatý Kůň | Pod_Hradem | S | Czech Republic | 49.37 N, 16.72 E | ~45-40 | (Nejman et al., 2017) |
| Zlatý Kůň | Moravský_Krumlov_IV | S | Czech Republic | 49.05 N, 16.41 E | ~43-42 | (Nerudová & Neruda, 2017) |
| Zlatý Kůň | Stranska_Skala_III-IIIc | IUP | Czech Republic | 49.11 N, 16.40 | ~45-40 | (Tostevin, 2003) |
| Zlatý Kůň | Brno-Bohunice | IUP | Czech Republic | ~49.12 N, 16.37 E | >45 | (Richter et al., 2008) |

**1.1 IUP-UP definition**

The definition of a techno-complex is based on the association among specific traits identified and recorded in archaeological assemblages (e.g. typology and technology of stone tools). Depending on the degree of magnification used to define a lithic assemblage, each techno-complex could be seen as more/less similar to others. A broad definition of techno-complexes could be compatible with models of population structure/movement, but this requires a high degree of simplification and the loss of finer-grained differences across the examined sites. On the other hand, a strict definition of a techno-complex could focus on the specificities of each assemblage, hampering the possibility of gaining a more general perspective. Strict definitions of lithic assemblages are in fact more compatible with scenarios of independent development (N. Zwyns & Lbova, 2019). In order to develop a model which considers a wide geographical and chronological scale as the one under consideration in this work, it is crucial to describe the techno-complexes choosing the key criteria that are both able to: i) maintain site-specificity; and ii) identify the general tendencies. The question is how the data are collected and compared and what meaning can be given to them concerning the question of the origin of technical innovations. New technologies could be related to demic movements, cultural diffusion/exchange without implying migration, or parallel convergence, i.e. the same response to common adaptive challenges. Distinguishing homoplasy from homology requires testing a hypothesis on the diversity of cultural traits and understanding it through a comprehensive framework of technological evolution(E. Boëda et al., 2013; Boyd & Richerson, 1988; Kuhn, 2019; Morgan et al., 2014; Peng et al., 2020).

The terms Initial Upper Paleolithic, Early Upper Paleolithic and Upper Paleolithic, are commonly used in the scientific literature with different meanings. They could have either a chronological connotation or a technological one, or both of them. This paper refers to the term Initial Upper Paleolithic (IUP) to indicate specific techno-complexes characterised by volumetric blade productions and Levallois reduction sequence(Kuhn, 2019; Kuhn & Zwyns, 2014) (Table S2). The term Upper Paleolithic (UP) is used to group the lithic industries characterised by the production of several standardised blades and bladelets often together with ornaments and bone tools (Table S2). The non-Mousterian and non-IUP technologies appeared during the Middle to Upper Palaeolithic transition, comprising Uluzzian(Benazzi et al., 2011; Collina et al., 2020; Marciani et al., 2020; Moroni et al., 2018; Peresani et al., 2019; Riel-Salvatore, 2009), Châtelperronian(M. Roussel et al., 2016; Morgan Roussel, 2013; Ruebens et al., 2015), Szeletian(Zsolt Mester, 2018; Neruda & Nerudová, 2019; Nerudová & Neruda, 2017) and Lincombian-Ranisian-Jermanowician (LRJ)(Flas, 2011; Kot et al., 2020; Krajcarz et al., 2018).

**Table S2. Main criteria adopted in this work to characterise Initial Upper Paleolithic (IUP) and Upper Paleolithic (UP) sites.**

| **Definition** | **IUP**<br><br>**Initial Upper Paleolithic** | **UP**<br><br>**Upper Paleolithic** |
|---|---|---|
| Lithic technology | The IUP is a blade-based production.<br><br>The reduction sequences use direct hard hammer percussion, platform faceting, and mostly flat-faced or semi-tournant cores. The blades and some cores resemble products of Levallois reduction sequence(Kuhn, 2019; Kuhn et al., 2009; Kuhn & Zwyns, 2014). Sites in Siberia and Mongolia are also characterised by “burin-cores” for producing small blades and the exploitation of the narrow face cores(Slavinsky et al., 2019; N. Zwyns et al., 2012; N. Zwyns & Lbova, 2019). | Blade and bladelet based industries usually used in complementary tools.<br><br>The broad category of UP includes a great diversity of techno-complexes which share the production of standardised bladelets and blades with a wide range of technical options (e.g. unidirectional debitage by prismatic core, carinated core, burin-cores, among others) and percussion techniques(Bataille, 2016; Falcucci, 2018; Kadowaki et al., 2021b; Kozłowski, 2015b; Le Brun-Ricalens et al., 2009; Moreau, 2012a; Teyssandier et al., 2010a; Zilhão et al., 2006). |
| Included techno-complex | Emiran(Kuhn et al., 1999), Bokerian(Leder, 2014, 2017), Bohunician(Yuri E. Demidenko et al., 2020; Richter et al., 2008; Škrdla, 2017), Bachokirian(Hublin et al., 2020). | Protoaurignacian and Aurignacian(Bataille, 2016; Y. E. Demidenko & Škrdla, 2017; Falcucci, 2018; Riel-Salvatore & Negrino, 2018; Sitlivy et al., 2012; Tafelmaier, 2017; Teyssandier et al., 2010b), Spitsynian(Usik et al., 2006; Vishnyatsky & Nehoroshev, 2004), Ahmarian(Barzilai et al., 2016; Goring-Morris & Belfer-Cohen, 2018; |


6

| | | |
|---|---|---|
| | | Kadowaki et al., 2015), Early Upper Paleolithic(Hoffecker, 2011; Kadowaki et al., 2021a), Gravettian(Dobrovolskaya et al., 2012; Klaric, 2013; Kozłowski, 2015a; Moreau, 2012b; Andey A. Sinitsyn, 2007). |
| Other material evidence | In some IUP assemblage is documented the use of personal ornament and formal bone tools e.g. in Levant(Kuhn et al., 2009), Bulgaria, Siberia, and Mongolia(Derevianko & Rybin, 2003; Kaifu et al., 2014; Kuhn & Zwyns, 2014; Rybin, 2014). | Habitual use (especially in the Gravettian) of portable art, graphic representations, musical instruments, and various bone tools(Conard, 2003; Conard et al., 2009; Zilhão et al., 2006). Impressive burials sites e.g. Sunghir(Trinkaus et al., 2014). |
| Chronology | Chronologically, the IUP is a long phenomenon comprised between approximately 50 kya and 35 kya (calibrated). Its stratigraphic position follows the Middle Paleolithic assemblages and is before UP assemblage(Zsolt Mester, 2018; Neruda & Nerudová, 2019; Nerudová & Neruda, 2017)s(Kuhn, 2019; Kuhn & Zwyns, 2014). | The various stages of the UP make their appearances approximately 42 kya. Their stratigraphic position follows the IUP, Uluzzian, Szeletian, Chatelperroian, LRS. |

### Supplementary Section 2: Genetic dataset

We downloaded the aligned sequences for the IUP Bacho Kiro individuals from the European Nucleotide Archive (accession number PRJEB39134), and merged files from the same individual. We clipped 3 bases off the ends of each read to avoid excess of aDNA damage using trimBam (https://genome.sph.umich.edu/wiki/BamUtil:_trimBam) and then generated pseudo-haploid calls with the genotype caller pileupCaller (https://github.com/stschiff/sequenceTools/tree/master/src-pileupCaller) using the majorityCall option that, for each locus, either chooses the allele supported by the highest number of reads or picks one randomly if the number is the same. Only positions with base quality and read quality higher than 30 were considered.
The processed Eigenstrat files of the Zlatý Kůň individual were kindly provided by Prof. Cosimo Posth and Dr. He Yu.
We merged the files with the 1240K “Allen Ancient DNA Resource” v44.3 database (https://reich.hms.harvard.edu/allen-ancient-dna-resource-aadr-downloadable-genotypes-present-day-and-ancient-dna-data)(Bergström et al., 2020; Skoglund et al., 2015), then kept only autosomal SNPs (1,150,639) using Plink 1.9(Chang et al., 2015) and then converted back to Eigenstrat format using Admixtools convertf(Patterson et al., 2012).

**Table S3: Individuals used in qpGraph modelling.** A green background indicates samples present in the "Allen Ancient DNA Resource" v44.3 database while an orange background indicates samples available from ENA.

| Version ID | Master ID | Publication | GroupID | qpGraph Pop ID |
|---|---|---|---|---|
| Chimp.REF | Chimp | Genome | Chimp.REF | Chimp |
| GoyetQ116-1_published | GoyetQ116-1 | FuNature2016 | Belgium_UP_GoyetQ116_1_published_all | GoyetQ116-1 |
| Kostenki14 | Kostenki14 | FuNature2016 | Russia_Kostenki14 | Kostenki14 |
| MA1.SG | MA1 | RaghavanNature2013 | Russia_MA1_HG.SG | MA1 (Mal'ta) |
| HGDP00449.SDG | HGDP00449 | BergstromScience2020 | Mbuti.SDG | Mbuti |
| HGDP00462.SDG | HGDP00462 | BergstromScience2020 | Mbuti.SDG | Mbuti |
| HGDP00463.SDG | HGDP00463 | BergstromScience2020 | Mbuti.SDG | Mbuti |
| HGDP00467.SDG | HGDP00467 | BergstromScience2020 | Mbuti.SDG | Mbuti |
| HGDP00474.SDG | HGDP00474 | BergstromScience2020 | Mbuti.SDG | Mbuti |
| HGDP00476.SDG | HGDP00476 | BergstromScience2020 | Mbuti.SDG | Mbuti |
| HGDP00478.SDG | HGDP00478 | BergstromScience2020 | Mbuti.SDG | Mbuti |
| HGDP00982.SDG | HGDP00982 | BergstromScience2020 | Mbuti.SDG | Mbuti |
| HGDP00984.SDG | HGDP00984 | BergstromScience2020 | Mbuti.SDG | Mbuti |
| HGDP01081.SDG | HGDP01081 | BergstromScience2020 | Mbuti.SDG | Mbuti |
| Oase1_d | Oase1 | FuNature2015 | Romania_Oase | Oase1 |
| B_Papuan-15.DG | HGDP00546 | PrueferNature2013 | Papuan.DG | Papuan |
| S_Papuan-1.DG | HGDP00550 | SkoglundNature2015 | Papuan.DG | Papuan |
| S_Papuan-10.DG | HGDP00553 | SkoglundNature2015 | Papuan.DG | Papuan |
| S_Papuan-11.DG | HGDP00555 | SkoglundNature2015 | Papuan.DG | Papuan |
| S_Papuan-12.DG | HGDP00556 | SkoglundNature2015 | Papuan.DG | Papuan |
| S_Papuan-13.DG | HGDP00552 | SkoglundNature2015 | Papuan.DG | Papuan |
| S_Papuan-14.DG | HGDP00554 | SkoglundNature2015 | Papuan.DG | Papuan |
| S_Papuan-2.DG | HGDP00540 | SkoglundNature2015 | Papuan.DG | Papuan |
| S_Papuan-3.DG | HGDP00541 | SkoglundNature2015 | Papuan.DG | Papuan |
| S_Papuan-4.DG | HGDP00543 | SkoglundNature2015 | Papuan.DG | Papuan |
| S_Papuan-5.DG | HGDP00545 | SkoglundNature2015 | Papuan.DG | Papuan |
| S_Papuan-6.DG | HGDP00547 | SkoglundNature2015 | Papuan.DG | Papuan |


8

| | | | | |
|---|---|---|---|---|
| S_Papuan-7.DG | HGDP00548 | SkoglundNature2015 | Papuan.DG | Papuan |
| S_Papuan-8.DG | HGDP00549 | SkoglundNature2015 | Papuan.DG | Papuan |
| S_Papuan-9.DG | HGDP00542 | SkoglundNature2015 | Papuan.DG | Papuan |
| baa001.SG | baa001 | SchlebuschScience2017 | South_Africa_1900BP.SG | South_Africa_2000BP |
| bab001.SG | bab001 | SchlebuschScience2017 | South_Africa_2000BP.SG | South_Africa_2000BP |
| I9028.SG | KhoesanLeipzigHunter | SkoglundCell2017 | South_Africa_2200BP.SG | South_Africa_2000BP |
| I9133.SG | UCT386 | SkoglundCell2017 | South_Africa_1900BP.SG | South_Africa_2000BP |
| Sunghir1.SG | Sunghir1 | SikoraScience2017 | Russia_Sunghir1.SG | Sunghir |
| Sunghir2.SG | Sunghir2 | SikoraScience2017 | Russia_Sunghir2.SG | Sunghir |
| Sunghir3.SG | Sunghir3 | SikoraScience2017 | Russia_Sunghir3.SG | Sunghir |
| Sunghir4.SG | Sunghir4 | SikoraScience2017 | Russia_Sunghir4.SG | Sunghir |
| Tianyuan | Tianyuan | YangCurrentBiology2017 | China_Tianyuan | Tianyuan |
| Ust_Ishim_published.DG | Ust_Ishim | FuNature2014 | Russia_Ust_Ishim_HG_published.DG | Ust_Ishim |
| Vindija_snpAD.DG | Vindija | Pruefer2017 | Vindija_Neanderthal.DG | Vindija_Neanderthal |
| Yana_old.SG | Yana1 | SikoraNature2019 | Russia_Yana_UP.SG | Yana |
| Yana_old2.SG | Yana2 | SikoraNature2019 | Russia_Yana_UP.SG | Yana |
| ZKU002 | ZKU002 | Pruefer2021 | ZlatyKun | ZlatyKun |
| BB7-240 | BB7-240 | Hajdinjak2021 | Bacho_Kiro | Bacho_Kiro |
| CC7-335 | CC7-335 | Hajdinjak2021 | Bacho_Kiro | Bacho_Kiro |
| F6-620 | F6-620 | Hajdinjak2021 | Bacho_Kiro | Bacho_Kiro |

### Supplementary Section 3: qpGraph analyses

We ran Admixtools qpGraph(Patterson et al., 2012), https://github.com/DReichLab/AdmixTools, qpGraph version: 7365) with the following parameters: outpop: NULL, blgsize: 0.05, lsqmode: YES, diag: .0001, useallsnps: NO, bigiter: 6, forcezmode: YES, initmix: 10000, precision: .0001, zthresh: 3.0, terse: NO, hires: YES.

#### 3.1 Base graph with Zlatý Kůň and Eurasian trifurcation

The position of Ust'Ishim with respect to Paleolithic Eastern and Western Eurasian is unclear and described as a near-trifurcation(Lipson & Reich, 2017; Yang et al., 2017) with D-statistics in the form of (Tianyuan, Sunghir, Ust'Ishim, Chimp), (Tianyuan, Ust'Ishim, Sunghir, Chimp) and (Sunghir, Ust'Ishim, Tianyuan, Chimp) all not significantly different from


9

zero (Table S4).

We started with the simple population tree proposed by Prufer and colleagues(Prüfer et al., 2021) (Fig 2c) with Zlatý Kůň as the most basal non African and placed Ust'Ishim either basal to the split between Western and Eastern Eurasians, as a sister of East Eurasians or as a sister of West Eurasians and found that all options are supported with the worst |Z-score| < 3. We used Tianyuan as "East Asian" and either Sunghir or Kostenki14 as "West Eurasian".

**3.2 Adding Bacho Kiro**

We then tried to add the IUP Bacho Kiro individuals in different positions of these graphs, downstream of the Zlatý Kůň split. We assumed Zlatý Kůň to be more basal than Bacho Kiro because of the following reasons:

- The Bacho Kiro individuals, set aside their higher Neanderthal admixture proportion, have been shown to share more alleles with East Asians compared to Europeans, while Zlatý Kůň is basal to the split between the two(Hajdinjak et al., 2021).
- f4 in the form (X, Zlatý Kůň, Bacho Kiro, Chimp) with X a later Eurasian individual including Ust'Ishim tend to be positive, even if not always significant (Table S4).

In order to allow the Neanderthal population that admixed with the ancestors of Bacho Kiro (likely living in Europe) to be different from the Neanderthal population that admixed with the ancestors of all non Africans shortly after the Out of Africa (likely in the Middle East) we modeled it as a different node closer to the Neanderthal from Vindija (Croatia); if it was indeed the same Neanderthal population the inferred drift between the two nodes would be 0. Additionally, in order to avoid our results to be driven by later population interaction between Eurasia and western Africa(Chen et al., 2020) and/or by the putative admixture of Mbuti pygmies with an archaic ghost hominin(Hammer et al., 2011; Hsieh et al., 2016; Lipson et al., 2020) we used four ancient South African hunter gatherers (Table S4) instead of Mbuti.

As a first attempt we placed Bacho Kiro basal to Ust'Ishim and all other Eurasians except Zlatý Kůň (Figure S1.A): the resulting graph has several f4 with |Z-scores| > 3: Bacho Kiro needs to be closer to Tianyuan (f2 = -3) and further from Sunghir (f2 = 2.6). A similar result is obtained when placing Ust'Ishim basal to Bacho Kiro and Bacho Kiro basal to the split between Europeans and East Asians, here represented by Tianyuan and Sunghir (Figure S1.B).

Rather than adding an admixture event from a Bacho Kiro related population to Tianyuan we tried to minimize admixture events and instead placed Bacho Kiro as a sister of Tianyuan hence leaving Ust'Ishim basal to the split Europe/East Asia. This returned a graph with the worst Z-score = -2.8 (Figure S1.C).

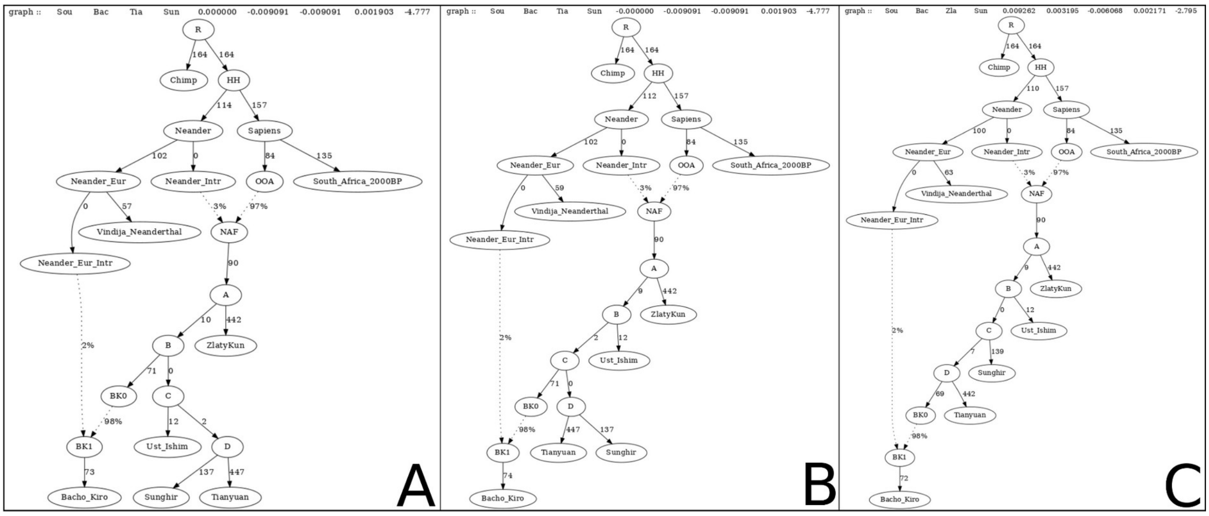


**Figure S1 Placement of Bacho Kiro within the tree proposed in Prufer et al. 2021.** When taking into account the excess Neanderthal contribution, Bacho Kiro still yields outlier f4s when placed soon after the split of Zlatý Kůň (**A**) or after the separation of Ust'Ishim (**B**). Placing Bacho Kiro as a sister of Tianyuan (**C**) yields instead no f4 outliers. (Graphs based on 421374 SNPs)

The placement of Bacho Kiro that we are proposing with respect to other Eurasians differs from the one proposed by Hajdinjak and colleagues(Hajdinjak et al., 2021) in their Figure S6.2. We speculate this may be due to the presence of Zlatý Kůň in our tree who may inform the genetic drift that characterizes a basal OoA landscape, or to the availability of Neanderthal from the early phases of our qpGraph construction (whose absence, by virtue of the additional Neanderthal ancestry compared to other non Africans, can have the effect of making Bacho Kiro appear more basal if not immediately accounted for)(Table S4).
For the sake of completeness we also tried Bacho Kiro as a sister of Sunghir (geographically it would make sense). The graph has several f4 |Z-scores| > 3 and is therefore rejected.

#### 3.3 Ust'Ishim as an early leaf of the IUP branch

To "populate" more the European branch we tried to add Kostenki14, the oldest "genetically European" individual sequenced to date (Figure S2.A – final score 12371). Since we noticed that the drift between nodes B and C was 0 and bearing in mind that D-statistics (Table S4). and previous graphs supported Sunghir/Tianyuan/Ust'Ishim to be a trifurcation, we tried to place Ust-Ishim either as a sister of Tianyuan/Bacho Kiro (Figure S2.B – score 12051) or as a sister of Kostenki14/Sunghir (Figure S2.C – score final 12173). All three graphs produce no f4 outliers and Ust'Ishim shares only one unit of drift with its sisters before branching off (hence showing a near-trifurcation). The tree where Ust'Ishim is a sister of Tianyuan/Bacho Kiro (Figure S2.B), however, has the lowest final score and so is the most supported.

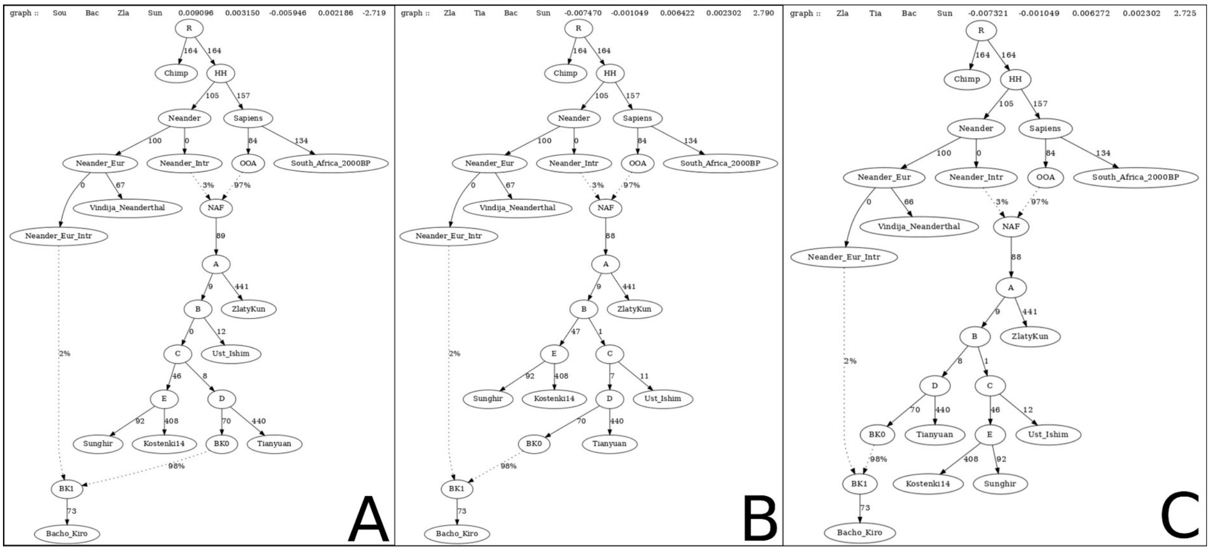


**Figure S2 Relationship of Ust'Ishim to other Eurasians.** After adding Kostenki14 as a crucial reference sample, we tested all possible positions of Ust'Ishim as either basal to East and West Eurasians (**A**), sister of Bacho Kiro and Tianyuan (**B**) and sister of Kostenki and Sunghir (**C**). All three graphs produce no f4 outliers and Ust'Ishim shares only one unit of drift with its sisters before branching off (hence showing a near-trifurcation). The tree where Ust'Ishim is a sister of Tianyuan/Bacho Kiro (Figure S2.B), however, has the lowest final score and so is the most supported. (Graphs based on 407081 SNPs)

#### 3.4 Oase1 could be a descendant of Bacho Kiro, remnant of the IUP movement

We then tried to place on the emerging tree of Figure S2.B the ~40 ky old Oase1 individual, recovered from a site less than 400 km away from Bacho Kiro cave which, like and more than the individuals from Bacho Kiro, has been shown to have a Neanderthal relative in its recent genealogy.

Since it has not been possible to determine a higher affinity of Oase1 for either East Asians or Europeans (retrospectively that might be caused by the combination of low number of SNPs and high Neanderthal admixture proportion), we started by making Oase1 a sister of Zlatý Kůň that later admixed with Neanderthals (Figure S3.A). The resulting graph showed several |Z scores| > 3, a way too high proportion of Neanderthal in Eurasians and a clear tendency for Oase1 and Bacho Kiro to share more drift (f2 = -6.3). We then modeled Oase1 as a sister of Bacho Kiro with an additional Neanderthal pulse and obtained no outlier f4 and a parsimonious placement for this elusive sample (Figure S3.B).

Our proposed graphs illustrate the claim made by Fu and colleagues(Fu et al., 2015) that Oase1 experienced an additional pulse of Neanderthal admixture between the one shared by all non Africans and the one that occurred 4-6 generations before it lived, and identify said event in the one occurred in the Bacho Kiro population or in a closely related one.
A similar graph where the node Oase0 splits from BK0 instead of BK1 (Figure S3.C) is also acceptable but does not explain this additional event of Neanderthal admixture.

To further support the placement of Oase1 and rule out the possibility that its attraction to Bacho Kiro is driven by the excess of Neanderthal ancestry they share, we proceeded to mask the most recent Neanderthal introgressed segments of Oase1 (we used the genomic


12

coordinates reported in Table S5.1 of Fu et al 2015(Fu et al., 2015)) and re-run the analysis: Oase1 holds its position and can be modeled without the last admixture event with Neandethal hence confirming its genuine connection with the Bacho Kiro population or a related one.

Finally, it is worth mentioning that while the low coverage, high contamination and high Neanderthal ancestry of Oase1 prevented the direct assessment of its closer relationship to either Western or Eastern Eurasians, an individual from the same site and with similar age (Oase2) showed a clearly higher affinity for East Asian and Native American populations than with Western Eurasians. The closest sample to Oase2 in outgroup f3 analyses, after Oase1, was reported to be Tianyuan (the individuals from Bacho Kiro cave were not available at the time of those analyses), supporting our claim of its placement in the "genetically East Asian" branch(Siska, 2019).

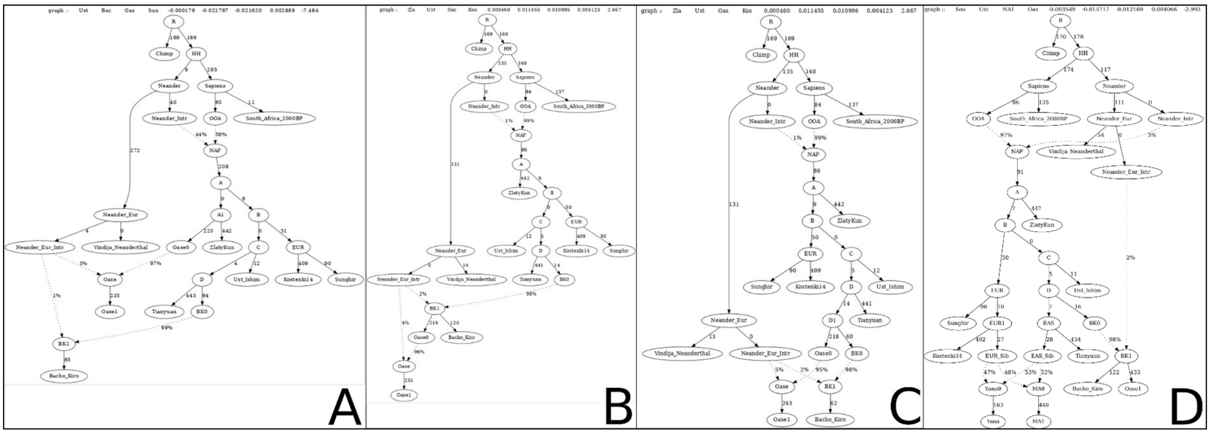


**Figure S3 Placement of Oase along the IUP tree.** After accounting for the reported recent pulse of Neanderthal admixture, Oase1 cannot be modelled as a simple basal lineage along the Eurasian tree (**A**) and is instead best fit as a descendant of the same population Bacho Kiro belonged to (**B**). This model provides an explanation for the putative three Neanderthal admixtures detected in Oase1 and is consistent with its younger date compared to Bacho Kiro and is therefore to be preferred over an equally fitting model (**C**) where Bacho Kiro and Oase1 have instead parallel evolutionary histories (graphs based on 95100 SNPs). When masking the very recent Neanderthal introgressed segments present in Oase1, this individual can be modeled without requiring the third and last admixture event with Neanderthals confirming its genuine attraction to the Bacho Kiro population (**D**). (Graphs A, B, C based on 95100 SNPs, graph D based on 66513 SNPs).

#### 3.5 Interaction between Zlatý Kůň and Bacho Kiro

Given the geographical proximity of the two sites we tested whether a contribution to the Bacho Kiro individuals by a population related to Zlatý Kůň is tolerated by our proposed model.

An admixture event contributing between 2% and 20% from a sister of Zlatý Kůň into the ancestors of Bacho Kiro, depending on the position of Ust'Ishim (Figure S4), yields no outliers. Although not rejected these graphs are all less parsimonious than the graph in Figure S2.B and might pick up a signal of shared drift between Bacho Kiro and Zlatý Kůň that is caused by the reduced time purifying selection had to drive out introgressed Neanderthal segments (Zlatý Kůň is the oldest sample and Bacho Kiro has recent Neanderthal introgression), rather than a genuine gene flow.


13

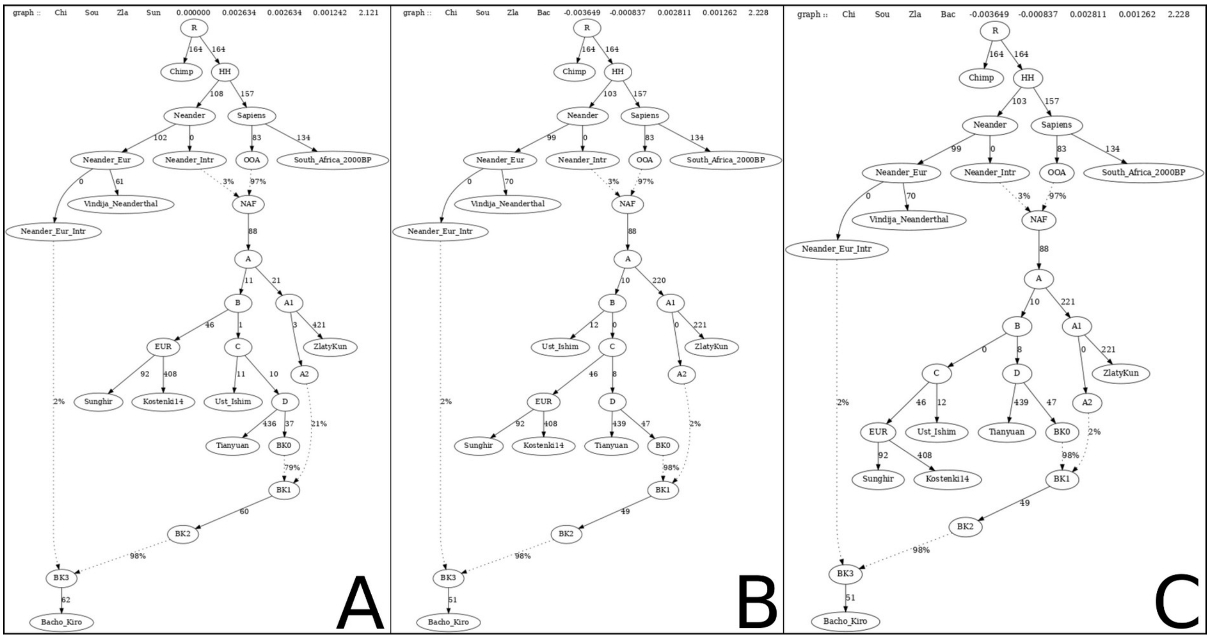


**Figure S4 Interaction between Zlatý Kůň and Bacho Kiro.** Given the geographic and partial chronological overlap of Zlatý Kůň and Bacho Kiro samples we allowed for a contribution of the former on the newly arrived IUP population. Such a contribution is acceptable, in varying amounts depending on the position of Ust'Ishim within the tree (**A,B,C**) but given the viability of the trees without this event we decided to exclude it from further analyses. (Graphs based on 407081 SNPs).

#### 3.6 Formation of Paleolithic Siberian ancestry profile

Paleolithic Siberian populations younger than 40 ky are consistently described as a mix of European and East Asian ancestries(Massilani et al., 2020; Raghavan et al., 2014; Sikora et al., 2019), although a comprehensive cultural or population dynamics to account for these admixture events is still lacking. Building on our dichotomy between Initial Upper Paleolithic (IUP) and Upper Paleolithic waves of expansions across Eurasia, we tried to model ancient Siberian individuals as a mixture of these waves into our best fitting parsimonious graph (Figure S2.B).

We started by creating a Siberian population that is ancestral to the ~32 ky old Yana individuals(Massilani et al., 2020; Raghavan et al., 2014; Sikora et al., 2019) as a mixture of the E and C nodes in the graph of Figure S2.B. The resulting graph has several f4 outliers and indicates that Yana may fit better as more closely related to Tianyuan (f2 = -6.1) and, to a lesser extent, Bacho Kiro (f2 = -1.9), suggesting that the node of origin of the East Asian component in Siberians needs to be moved further down along that branch. With the East Asian component of Siberians originating from node D the graph still has several outliers and Yana needs to share more drift with Tianyuan (f2 = -3.6), so after taking that into account we obtained a graph that is not rejected (Figure S4.A).
To better define the Siberian populations we included the ~24 kya Mal'ta (MA1) individual(Fu et al., 2015; Raghavan et al., 2014), we started by considering it as a sister of Yana (Figure S4.B) and obtained two |Z-scores| ~3,1 (Chimp, South_Africa_2000BP, ZlatyKun, Yana and ZlatyKun, Tianyuan, Sunghir, MA1). Since qpGraph performs f-statistic analyses on a subset of SNPs that is shared by all the individuals in a model, this might sometimes severely limit the number of SNPs used in a test, especially when ancient individuals are included. We

tested whether the two outliers remain significantly different from zero when including all SNPs available to each quadruple and found they do not (Table S4), deeming this outcome as more robust.
Finally, in order to allow MA1 and Yana to have different proportions of East and West Eurasian components, we added an EUR_Sib node and parallelized the admixture events of Yana and MA1. The resulting graph (Figure S5.C) has two Dstat Z-scores (Chimp, South_Africa_2000BP, ZlatyKun, Yana and ZlatyKun, Tianyuan, Sunghir, MA1) higher than 3 in absolute value (the same two that were present earlier, with Z = 3.1 and -3.0 respectively), however both of them result not significantly different from zero when computing the D-statistic using the all the available SNPs (Table S4) .

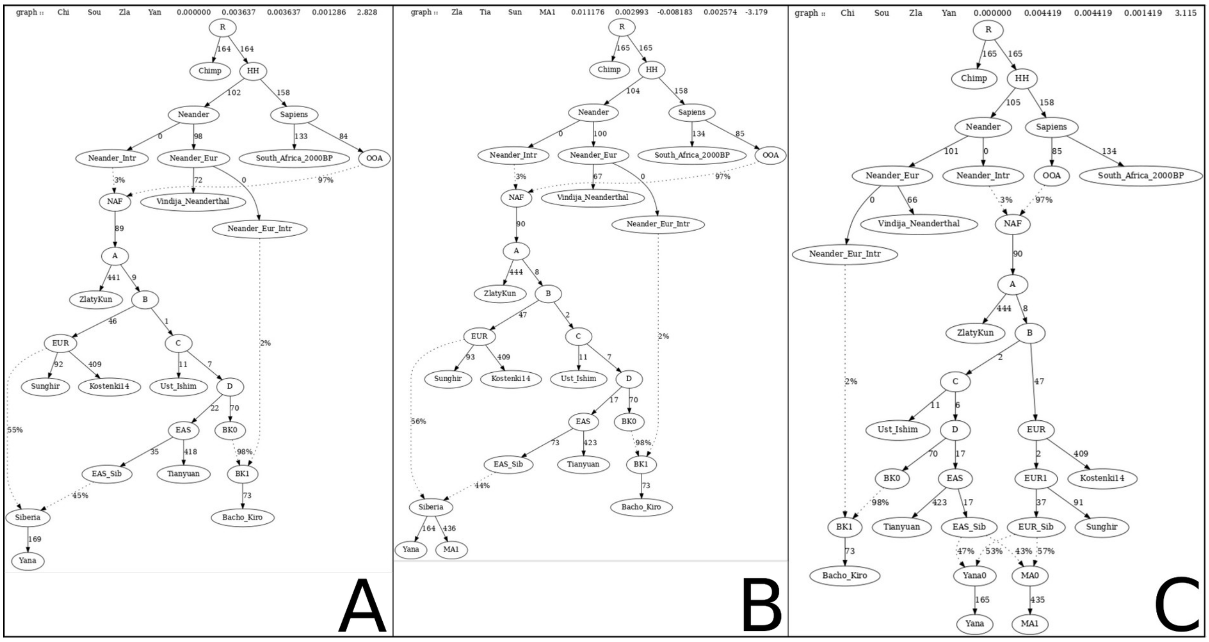


**Figure S5 Adding Paleolithic Siberians to the qpGraph of Figure S2.B.** We first placed Yana1 on the graph as a putative mixture of West Eurasian (EUR) and East Asian (EAS) components (**A**) and, after having obtained no outliers we proceeded with adding Mal'ta as a sister leaf (**B**). To allow for the two Siberian samples to derive independent fractions of each ancestry we then relaxed the proposed model to obtain (**C**). (Graph A based on 412809 SNPs, graphs B and C based on 302476 SNPs).

#### 3.7 Adding GoyetQ116-1 to the emerging picture of IUP and UP interaction

The ~35000 years old GoyetQ116-1 individual from Belgium, although closer to Europeans than to East Asians, shares more alleles with the latter compared to other contemporary Europeans(Fu et al., 2016) and it has recently been described as a recipient of gene flow from a population related to Bacho Kiro(Hajdinjak et al., 2021).

We started from Figure S5.C and modeled Goyet as mixture between the Kostenki14/Sunghir branch and the Bacho Kiro branch (Figure S6.A); however doing so shows that Goyet needs to share more drift with Tianyuan (f2 = -2.4, plus several f4 outliers) highlighting how its East Asian genetic component is more variegated than the one found in Bacho Kiro or Tianyuan alone.
When adding to Goyet a contribution from both the Bacho Kiro and Tianyuan branches we obtained only two minor outliers (Chimp, South_Africa_2000BP, ZlatyKun, Yana = 3.4 and South_Africa_2000BP, ZlatyKun, ZlatyKun, Sunghir = -3.3), the first of which results non


15

significant when computed using all SNPs (Table S4). The results are comparable whether the two genetically East Asian components admix with each other first, and then with the European one (Figure S6.B), or vice versa (Figure S6.C).

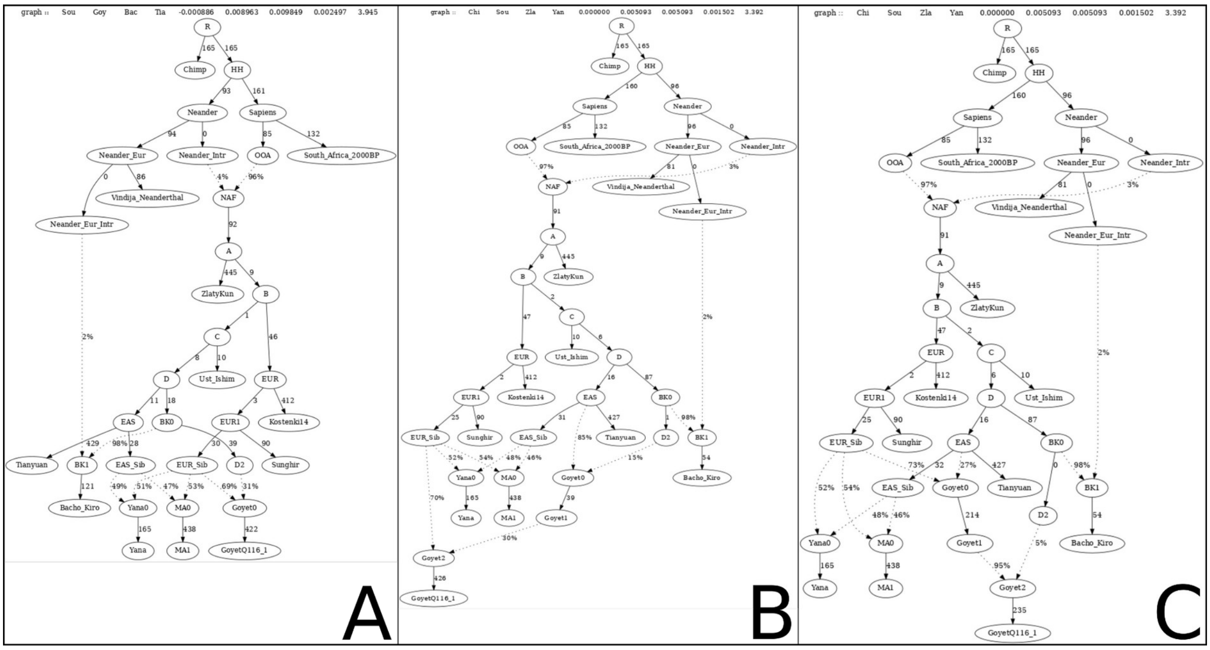


**Figure S6 Goyet Q116-1 as a mixture of UP and IUP lineages.** Modelling the East Asian component found in Goyet Q116-1 as a simple interaction between UP and the Bacho Kiro branch (**A**) yields an unexplained attraction between Goyet Q116-1 and Tianyuan. Modelling the IUP component as a mixture of Bacho Kiro- and Tianiyuan-related lineages before (**B**) or after (**C**) the arrival of the UP lineages provides a better fit to the data. (Graphs based on 256790 SNPs).

#### 3.8 Peopling of Oceania

The position of Oceanian populations with respect to the East and West Eurasian is still unclear, with Oceanians often seen as either an earlier split (Choin et al., 2021; Malaspinas et al., 2016) or a sister population to East Asians(Mallick et al., 2016; Wall, 2017). Given the relatively simple population tree needed to explain the post OoA Eurasian population history using aDNA samples available to date, and benefiting from the basal position of Zlatý Kůň, we tried to model Oceanian populations (modern Papuans) within our hypothesized model. We avoided including the sampled Denisova aDNA(Meyer et al., 2012a) within the population tree to eliminate attractions from yet uncharacterized, deep splits along the hominin branch, and opted for a surrogate ghost split along the archaic human lineage to account for the documented Denisova presence in Papuans(Choin et al., 2021; Malaspinas et al., 2016; Meyer et al., 2012b; Reich et al., 2011).

We started from Graph S2.B and tried to have Papuans split either before or after the split of the Zlatý Kůň lineage; in both cases the graph was rejected as papuans needed to share more drift with Tianyuan(f2: -11 and -8). For this reason we then modeled them as a sister group of Tianyuan and obtained a graph whose only two outliers (South_Africa_2000BP, Bacho Kiro, Zlatý Kůň, Papuan = -3.4 and Zlatý Kůň, Papuan, Bacho Kiro, Sunghir = 3.0) resulted non significant when taking all SNPs into account (Figure 1.B, Figure S7.A). Subsequently we tried to model Papuans as a mixture of a Tianyuan sister population and a more basal lineage; the contribution from said lineage decreases the more that split is moved backward: it is 96% when splitting just before the split Tianyuan/Bacho Kiro (Figure S7.B), 53% when splitting before Ust'Ishim (Figure S7.C), 40% when splitting after Zlatý Kůň but before any other Eurasian branching (Figure S7.D) and only 2% when splitting before Zlatý Kůň (Figure S7.E). Finally, we decided to test whether a small contribution to Papuans from a population of AMH that left Africa before the 70-60 kya Out of Africa (xOoA) (Pagani et al., 2016) is rejected by the model (Figure S7.F). A 2% contribution from said population to modern papuans is not rejected since the only f4 outlier(Zlatý Kůň, Papuans, Bacho Kiro, Sunghir = 3.0) is ~0 when including all available SNPs (Table S4).

All the acceptable solutions for the placement of Papuans within the broader OoA tree confirm Zlatý Kůň as the most basal human genome among the ones ever found Out of Africa. It is therefore reasonable to describe Oceanians as either an almost even mixture between East Asians and a lineage basal to West and East Asians occurred sometimes between 45 and 38kya, or as a sister lineage of East Asians with or without a minor basal OoA or xOoA contribution.


17

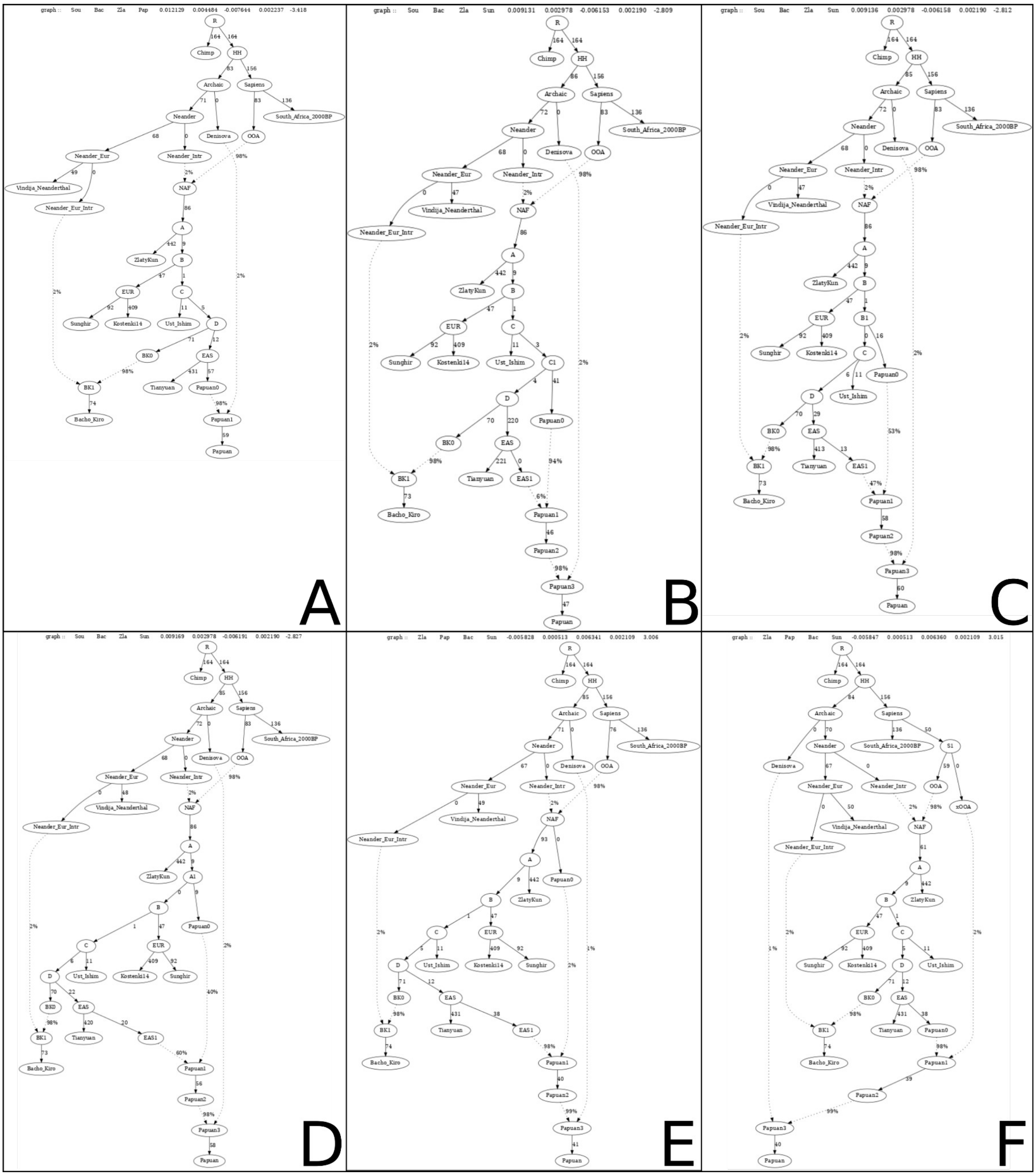


**Figure S7 The place of Papuans within the out of Africa graph.** Modern Papuans can be invariably described as a sister population of Tianyuan (**A**) with or without the addition of a more basal, *H. sapiens* lineage. The position of this deeper lineage along the population model influences the genetic contribution it gives to Papuans: 94% if placed within the IUP branch after Ust'Ishim (**B**), 53% before Ust'Ishim (**C**), 4% before the separation of IUP and UP branches (**D**) and 2% either as the most basal OoA branch (**E**) or even as an extinct, extra OoA (xOoA) (**F**). (Graphs based on 417400 SNPs).


18

**Table S4: Relevant f4 tests.** The first seven columns report population and values as outputted by Admixtools. The eight column reports the Supplementary Section where a given f4 is mentioned.

| Pop1 | Pop2 | Pop3 | Pop4 | D | Z | SNPs | Supp. Section |
|---|---|---|---|---|---|---|---|
| Ust_Ishim | Kostenki14 | Tianyuan | Chimp | 0.0006 | 0.087 | 817238 | 3.1 |
| Ust_Ishim | Sunghir | Tianyuan | Chimp | -0.0023 | -0.34 | 847217 | 3.1 |
| Tianyuan | Kostenki14 | Ust_Ishim | Chimp | -0.0037 | -0.523 | 817238 | 3.1 |
| Tianyuan | Sunghir | Ust_Ishim | Chimp | 0.0024 | 0.386 | 847217 | 3.1 |
| Ust_Ishim | Tianyuan | Kostenki14 | Chimp | 0.0043 | 0.588 | 817238 | 3.1 |
| Ust_Ishim | Tianyuan | Sunghir | Chimp | -0.0047 | -0.758 | 847217 | 3.1 |
| Bacho_Kiro | Kostenki14 | Ust_Ishim | Chimp | -0.0263 | -**4.088** | 997593 | 3.2 |
| Bacho_Kiro | Sunghir | Ust_Ishim | Chimp | -0.0178 | **-3.261** | 1052955 | 3.2 |
| Bacho_Kiro | Tianyuan | Ust_Ishim | Chimp | -0.0207 | **-3.223** | 841030 | 3.2 |
| Tianyuan | ZlatyKun | Bacho_Kiro | Chimp | 0.0364 | **5.142** | 554809 | 3.2 |
| Kostenki14 | ZlatyKun | Bacho_Kiro | Chimp | 0.0131 | 1.878 | 594315 | 3.2 |
| Sunghir | ZlatyKun | Bacho_Kiro | Chimp | 0.0138 | 2.199 | 620949 | 3.2 |
| Ust_Ishim | ZlatyKun | Bacho_Kiro | Chimp | 0.0192 | 2.762 | 619549 | 3.2 |
| ZlatyKun | Tianyuan | Sunghir | MA1 | 0.0085 | 1.112 | 422908 | 3.6 |
| Chimp | South_Africa_2000BP | ZlatyKun | Yana | 0.0122 | 2.864 | 625628 | 3.6, 3.7 |
| South_Africa_2000BP | Bacho_Kiro | ZlatyKun | Papuan | 0.0115 | 1.975 | 636119 | 3.8 |
| ZlatyKun | Papuan | Bacho_Kiro | Sunghir | 0.0003 | 0.046 | 636130 | 3.8 |
| Kostenki14 | ZlatyKun | Papuan | Chimp | 0.029 | **4.028** | 587236 | 3.8 |
| Sunghir | ZlatyKun | Papuan | Chimp | 0.027 | **4.452** | 614406 | 3.8 |
| Ust_Ishim | ZlatyKun | Papuan | Chimp | 0.0363 | **5.213** | 613032 | 3.8 |
| Tianyuan | ZlatyKun | Papuan | Chimp | 0.0736 | **10.746** | 547447 | 3.8 |


19

### Supplementary Section 4: Paleomaps plotting

To plot the paleomaps, we used R (version 4.0.5 - “Shake and Throw”), the GUI RStudio (version 1.4.1103 - “Wax Begonia”) and the package “oce” (https://dankelley.github.io/oce/). We downloaded the alti-bathymetric maps from the ETOPO1 dataset (Amante & Eakins, 2009) and used the inferred sea level values at given times in the past from the “Global 1Ma Temperature, Sea Level, and Ice Volume Reconstructions” dataset (Bintanja et al., 2005) available from the NOAA Paleoclimatology Program.
We plotted the maps applying Lambert conformal conic projections ("+proj=lcc +lat_1=25 +lat_2=50 +lon_0=97" in PROJ.4 format).

Bergström, A., McCarthy, S. A., Hui, R., Almarri, M. A., Ayub, Q., Danecek, P., Chen, Y., Felkel, S., Hallast, P., Kamm, J., Blanché, H., Deleuze, J.-F., Cann, H., Mallick, S.,

 20

Reich, D., Sandhu, M. S., Skoglund, P., Scally, A., Xue, Y., … Tyler-Smith, C. (2020). Insights into human genetic variation and population history from 929 diverse genomes. *Science*, *367*(6484). https://doi.org/10.1126/science.aay5012

Bintanja, R., van de Wal, R. S. W., & Oerlemans, J. (2005). Modelled atmospheric temperatures and global sea levels over the past million years. *Nature*, *437*(7055), 125–128.

Boëda, E., & Bonilauri, S. (2006). THE FIRST BLADELET PRODUCTION 40,000 YEARS AGO. *L'Anthropologie*, *44*(1), 75–92.

Boëda, E., Hou, Y. M., Forestier, H., Sarel, J., & Wang, H. M. (2013). Levallois and non-Levallois blade production at Shuidonggou in Ningxia, North China. *Quaternary International: The Journal of the International Union for Quaternary Research*, *295*, 191–203.

Bosch, M. D., Mannino, M. A., Prendergast, A. L., O'Connell, T. C., Demarchi, B., Taylor, S. M., Niven, L., van der Plicht, J., & Hublin, J.-J. (2015). New chronology for Ksâr 'Akil (Lebanon) supports Levantine route of modern human dispersal into Europe. *Proceedings of the National Academy of Sciences of the United States of America*, *112*(25), 7683–7688.

Boyd, R., & Richerson, P. J. (1988). *Culture and the Evolutionary Process*. University of Chicago Press.

Chang, C. C., Chow, C. C., Tellier, L. C., Vattikuti, S., Purcell, S. M., & Lee, J. J. (2015). Second-generation PLINK: rising to the challenge of larger and richer datasets. *GigaScience*, *4*, 7.

Chen, L., Wolf, A. B., Fu, W., Li, L., & Akey, J. M. (2020). Identifying and Interpreting Apparent Neanderthal Ancestry in African Individuals. *Cell*, *180*(4), 677–687.e16.

Choin, J., Mendoza-Revilla, J., Arauna, L. R., Cuadros-Espinoza, S., Cassar, O., Larena, M., Ko, A. M.-S., Harmant, C., Laurent, R., Verdu, P., Laval, G., Boland, A., Olaso, R., Deleuze, J.-F., Valentin, F., Ko, Y.-C., Jakobsson, M., Gessain, A., Excoffier, L., … Quintana-Murci, L. (2021). Genomic insights into population history and biological

 21

adaptation in Oceania. *Nature*, *592*(7855), 583–589.

Collina, C., Marciani, G., Martini, I., Donadio, C., Repola, L., Bortolini, E., Arrighi, S., Badino, F., Figus, C., Lugli, F., Oxilia, G., Romandini, M., Silvestrini, S., Piperno, M., & Benazzi, S. (2020). Refining the Uluzzian through a new lithic assemblage from Roccia San Sebastiano (Mondragone, southern Italy). In *Quaternary International* (Vol. 551, pp. 150–168). https://doi.org/10.1016/j.quaint.2020.03.056

Conard, N. J. (2003). Palaeolithic ivory sculptures from southwestern Germany and the origins of figurative art. *Nature*, *426*(6968), 830–832.

Conard, N. J., Malina, M., & Münzel, S. C. (2009). New flutes document the earliest musical tradition in southwestern Germany. *Nature*, *460*(7256), 737–740.

Demidenko, Y. E., & Škrdla, P. (2017). Aurignacian in Moravia New geochronological, lithic and settlement data. *Památky Archeologické*. http://search.ebscohost.com/login.aspx?direct=true&profile=ehost&scope=site&authtype=crawler&jrnl=00310506&AN=136138007&h=et6bISwXH0wKzdZ%2BYStr1w3WWX1C1u%2B7YNG%2FIjbjEJjQsRuL%2FHCXFatMGly9ew5GfXMK84vFUpUNdc7oDo9%2F2g%3D%3D&crl=c

Demidenko, Y. E., Škrdla, P., & Rychtaříková, T. (2020). Initial Upper Paleolithic bladelet production: Bladelets in Moravian Bohunician. In *Přehled výzkumů* (pp. 21–29). https://doi.org/10.47382/pv0611-02

Derevianko, A. P., & Rybin, E. P. (2003). The earliest representations of symbolic behavior by Paleolithic humans in the Altai Mountains. *Archaeology, Ethnology and Anthropology of Eurasia*, *3*, 27–50.

Derevianko, A. P., Rybin, E. P., Gladyshev, S. A., Gunchinsuren, B., Tsybankov, A. A., & Olsen, J. W. (2013). Early Upper Paleolithic Stone Tool Technologies of Northern Mongolia: The Case of Tolbor-4 and Tolbor-15*. *Archaeology, Ethnology and Anthropology of Eurasia*, *41*(4), 21–37.

Dobrovolskaya, M., Richards, M.-P., & Trinkaus, E. (2012). Direct radiocarbon dates for the Mid Upper Paleolithic (eastern Gravettian) burials from Sunghir, Russia. *Bulletins et Memoires de La Societe D’anthropologie de Paris*, *24*(1-2), 96–102.

 22

Falcucci, A. (2018). Towards a renewed definition of the Protoaurignacian. *Mitteilungen Der Gesellschaft Für Urgeschichte*, *27*, 87–130.

Flas, D. (2011). The Middle to Upper Paleolithic transition in Northern Europe: the Lincombian-Ranisian-Jerzmanowician and the issue of acculturation of the last Neanderthals. In *World Archaeology* (Vol. 43, Issue 4, pp. 605–627). https://doi.org/10.1080/00438243.2011.624725

Fu, Q., Hajdinjak, M., Moldovan, O. T., Constantin, S., Mallick, S., Skoglund, P., Patterson, N., Rohland, N., Lazaridis, I., Nickel, B., Viola, B., Prüfer, K., Meyer, M., Kelso, J., Reich, D., & Pääbo, S. (2015). An early modern human from Romania with a recent Neanderthal ancestor. *Nature*, *524*(7564), 216–219.

Fu, Q., Posth, C., Hajdinjak, M., Petr, M., Mallick, S., Fernandes, D., Furtwängler, A., Haak, W., Meyer, M., Mittnik, A., Nickel, B., Peltzer, A., Rohland, N., Slon, V., Talamo, S., Lazaridis, I., Lipson, M., Mathieson, I., Schiffels, S., … Reich, D. (2016). The genetic history of Ice Age Europe. *Nature*, *534*(7606), 200–205.

Goring-Morris, N., & Belfer-Cohen, A. (2018). The Ahmarian in the Context of the Earlier Upper Palaeolithic in the Near East. In Y. Nishiaki & T. Akazawa (Eds.), *The Middle and Upper Paleolithic Archeology of the Levant and Beyond* (pp. 87–104). Springer Singapore.

Hajdinjak, M., Mafessoni, F., Skov, L., Vernot, B., Hübner, A., Fu, Q., Essel, E., Nagel, S., Nickel, B., Richter, J., Moldovan, O. T., Constantin, S., Endarova, E., Zahariev, N., Spasov, R., Welker, F., Smith, G. M., Sinet-Mathiot, V., Paskulin, L., … Pääbo, S. (2021). Initial Upper Palaeolithic humans in Europe had recent Neanderthal ancestry. *Nature*, *592*(7853), 253–257.

Hammer, M. F., Woerner, A. E., Mendez, F. L., Watkins, J. C., & Wall, J. D. (2011). Genetic evidence for archaic admixture in Africa. *Proceedings of the National Academy of Sciences of the United States of America*, *108*(37), 15123–15128.

Hauck, T., Rethemeyer, J., & Rentzel, P. (2016). Neanderthals or early modern humans? A revised 14C chronology and geoarchaeological study of the Szeletian sequence in

Szeleta Cave (Kom. Borsod-Abaúj …. *Archäologisches*. https://edoc.unibas.ch/52894/

Hoffecker, J. F. (2011). The early upper Paleolithic of eastern Europe reconsidered. *Evolutionary Anthropology*, *20*(1), 24–39.

Hoffecker, J. F., Holliday, V. T., Anikovich, M. V., Dudin, A. E., Platonova, N. I., Popov, V. V., Levkovskaya, G. M., Kuz'mina, I. E., Syromyatnikova, E. V., Burova, N. D., Goldberg, P., Macphail, R. I., Forman, S. L., Carter, B. J., & Crawford, L. J. (2016). Kostenki 1 and the early Upper Paleolithic of Eastern Europe. *Journal of Archaeological Science: Reports*, *5*, 307–326.

Hsieh, P., Woerner, A. E., Wall, J. D., Lachance, J., Tishkoff, S. A., Gutenkunst, R. N., & Hammer, M. F. (2016). Model-based analyses of whole-genome data reveal a complex evolutionary history involving archaic introgression in Central African Pygmies. *Genome Research*, *26*(3), 291–300.

Hublin, J.-J., Sirakov, N., Aldeias, V., Bailey, S., Bard, E., Delvigne, V., Endarova, E., Fagault, Y., Fewlass, H., Hajdinjak, M., Kromer, B., Krumov, I., Marreiros, J., Martisius, N. L., Paskulin, L., Sinet-Mathiot, V., Meyer, M., Pääbo, S., Popov, V., … Tsanova, T. (2020). Initial Upper Palaeolithic Homo sapiens from Bacho Kiro Cave, Bulgaria. *Nature*, *581*(7808), 299–302.

Kadowaki, S., Omori, T., & Nishiaki, Y. (2015). Variability in Early Ahmarian lithic technology and its implications for the model of a Levantine origin of the Protoaurignacian. *Journal of Human Evolution*, *82*, 67–87.

Kadowaki, S., Suga, E., & Henry, D. O. (2021a). Frequency and production technology of bladelets in Late Middle Paleolithic, Initial Upper Paleolithic, and Early Upper Paleolithic (Ahmarian) assemblages in Jebel Qalkha, Southern Jordan. In *Quaternary International*. https://doi.org/10.1016/j.quaint.2021.03.012

Kadowaki, S., Suga, E., & Henry, D. O. (2021b). Frequency and production technology of bladelets in Late Middle Paleolithic, Initial Upper Paleolithic, and Early Upper Paleolithic (Ahmarian) assemblages in Jebel Qalkha, Southern Jordan. *Quaternary International: The Journal of the International Union for Quaternary Research*.



24

https://doi.org/10.1016/j.quaint.2021.03.012

Kaifu, Y., Izuho, M., Goebel, T., Sato, H., & Ono, A. (2014). *Emergence and Diversity of Modern Human Behavior in Paleolithic Asia*. Texas A&M University Press.

Khenzykhenova, F., Lipnina, E., Danukalova, G., Shchetnikov, A., Osipova, E., Semenei, E., Tumurov, E., & Lokhov, D. (2019). The area surrounding the world-famous geoarchaeological site Mal'ta (Baikal Siberia): New data on the chronology, archaeology, and fauna. *Quaternary International: The Journal of the International Union for Quaternary Research*, *509*, 17–29.

Klaric, L. (2013). Faciès lithiques et chronologie du Gravettien du sud du Bassin parisien et de sa marge sud-occidentale. *Pierre Bodu, Lucy Chehmana, Laurent Klaric, Ludovic Mevel, Sylvain Soriano éds*, *56*, 61–88.

Kot, M., Krajcarz, M. T., Hoyo, M. M., Gryczewska, N., Wojenka, M., Pyżewicz, K., Sinet-Mathiot, V., Diakowski, M., Fedorowicz, S., Gąsiorowski, M., Marciszak, A., & Mackiewicz, P. (2020). *Chronostratigraphy of Jerzmanowician. New data from Koziarnia Cave, Poland*. https://doi.org/10.1101/2020.04.29.067967

Kozłowski, J. K. (2015a). The origin of the gravettian. *Quaternary International: The Journal of the International Union for Quaternary Research*, *359-360*, 3–18.

Kozłowski, J. K. (2015b). The origin of the Gravettian. *Quaternary International: The Journal of the International Union for Quaternary Research*, *359-360*, 3–18.

Krajcarz, M. T., Krajcarz, M., Ginter, B., Goslar, T., & Wojtal, P. (2018). Towards a chronology of the jerzmanowician-a new series of radiocarbon dates from nietoperzowa cave (Poland). *Archaeometry*, *60*(2), 383–401.

Kuhn, S. L. (2004). Upper Paleolithic raw material economies at Üçağızlı cave, Turkey. *Journal of Anthropological Archaeology*, *23*(4), 431–448.

Kuhn, S. L. (2019). Initial Upper Paleolithic: A (near) global problem and a global opportunity. *Archaeological Research in Asia*, *17*, 2–8.

Kuhn, S. L., Stiner, M. C., & Güleç, E. (1999). Initial Upper Palaeolithic in south-central Turkey and its regional context: a preliminary report. *Antiquity*, *73*(281), 505–517.


25

Kuhn, S. L., Stiner, M. C., Güleç, E., Ozer, I., Yilmaz, H., Baykara, I., Açikkol, A., Goldberg, P., Molina, K. M., Unay, E., & Suata-Alpaslan, F. (2009). The early Upper Paleolithic occupations at Uçağizli Cave (Hatay, Turkey). *Journal of Human Evolution*, *56*(2), 87–113.

Kuhn, S. L., & Zwyns, N. (2014). Rethinking the initial Upper Paleolithic. In *Quaternary International* (Vol. 347, pp. 29–38). https://doi.org/10.1016/j.quaint.2014.05.040

Lbova, L. (2019). *Pigments on Upper Palaeolithic mobile art. Spectral analysis of figurines from Mal'ta culture (Siberia)*. HUGO OBERMAIER-GESELLSCHAFT für Erforschung des Eiszeitalters und der Steinzeit e.V. https://doi.org/10.7485/QU66_8

Le Brun-Ricalens, F. L., Bordes, J.-G., & Eizenberg, L. (2009). A crossed-glance between southern European and Middle-Near Eastern early Upper Palaeolithic lithic technocomplexes. Existing models, new perspectives. In *The Mediterranean from 50,000 to 25,000 bp: Turning Points and New Directions.* (Vol. 66, pp. 11–33).

Leder, D. (2014). *Technological and typological change at the Middle to Upper Palaeolithic Boundary in Lebanon*.

Leder, D. (2017). Core reduction strategies at the Initial Upper Palaeolithic sites of Ksar Akil and Abou Halka in Lebanon. *Lithics, The Journal of the Lithic Studies Society.* https://www.semanticscholar.org/paper/b3ae3d24daa6256817e3f70d774432699d38e22b

Li, F., Kuhn, S. L., Bar-Yosef, O., Chen, F.-Y., Peng, F., & Gao, X. (2019). History, Chronology and Techno-Typology of the Upper Paleolithic Sequence in the Shuidonggou Area, Northern China. *Journal of World Prehistory*, *32*(2), 111–141.

Lipson, M., & Reich, D. (2017). A Working Model of the Deep Relationships of Diverse Modern Human Genetic Lineages Outside of Africa. *Molecular Biology and Evolution*, *34*(4), 889–902.

Lipson, M., Ribot, I., Mallick, S., Rohland, N., Olalde, I., Adamski, N., Broomandkhoshbacht, N., Lawson, A. M., López, S., Oppenheimer, J., Stewardson, K., Asombang, R. N., Bocherens, H., Bradman, N., Culleton, B. J., Cornelissen, E., Crevecoeur, I., de Maret,


26

P., Fomine, F. L. M., … Reich, D. (2020). Ancient West African foragers in the context of African population history. *Nature*, *577*(7792), 665–670.

Malaspinas, A.-S., Westaway, M. C., Muller, C., Sousa, V. C., Lao, O., Alves, I., Bergström, A., Athanasiadis, G., Cheng, J. Y., Crawford, J. E., Heupink, T. H., Macholdt, E., Peischl, S., Rasmussen, S., Schiffels, S., Subramanian, S., Wright, J. L., Albrechtsen, A., Barbieri, C., … Willerslev, E. (2016). A genomic history of Aboriginal Australia. *Nature*, *538*(7624), 207–214.

Mallick, S., Li, H., Lipson, M., Mathieson, I., Gymrek, M., Racimo, F., Zhao, M., Chennagiri, N., Nordenfelt, S., Tandon, A., Skoglund, P., Lazaridis, I., Sankararaman, S., Fu, Q., Rohland, N., Renaud, G., Erlich, Y., Willems, T., Gallo, C., … Reich, D. (2016). The Simons Genome Diversity Project: 300 genomes from 142 diverse populations. *Nature*, *538*(7624), 201–206.

Marciani, G., Ronchitelli, A., Arrighi, S., Badino, F., Bortolini, E., Boscato, P., Boschin, F., Crezzini, J., Delpiano, D., Falcucci, A., Figus, C., Lugli, F., Oxilia, G., Romandini, M., Riel-Salvatore, J., Negrino, F., Peresani, M., Spinapolice, E. E., Moroni, A., & Benazzi, S. (2020). Lithic techno-complexes in Italy from 50 to 39 thousand years BP: An overview of lithic technological changes across the Middle-Upper Palaeolithic boundary. In *Quaternary International* (Vol. 551, pp. 123–149). https://doi.org/10.1016/j.quaint.2019.11.005

Marks, A. E., & Kaufman, D. (1983). Boker Tachtit: the artifacts. *Prehistory and Paleoenvironments in the Central Negev, Israel*, *3*, 69–126.

Massilani, D., Skov, L., Hajdinjak, M., Gunchinsuren, B., Tseveendorj, D., Yi, S., Lee, J., Nagel, S., Nickel, B., Devièse, T., Higham, T., Meyer, M., Kelso, J., Peter, B. M., & Pääbo, S. (2020). Denisovan ancestry and population history of early East Asians. *Science*, *370*(6516), 579–583.

Mester, Z. (2010). Technological analysis of Szeletian bifacial points from Szeleta Cave (Hungary). *Human Evolution*, *25*(1-2), 107–123.

Mester, Z. (2018). The problems of the Szeletian as seen from Hungary. In *Recherches*

*Archéologique Nouvelle Serie* (Vol. 9, pp. 19–48). https://doi.org/10.33547/rechacrac.ns9.02

Meyer, M., Kircher, M., Gansauge, M.-T., Li, H., Racimo, F., Mallick, S., Schraiber, J. G., Jay, F., Prüfer, K., de Filippo, C., Sudmant, P. H., Alkan, C., Fu, Q., Do, R., Rohland, N., Tandon, A., Siebauer, M., Green, R. E., Bryc, K., … Pääbo, S. (2012a). A high-coverage genome sequence from an archaic Denisovan individual. *Science*, *338*(6104), 222–226.

Meyer, M., Kircher, M., Gansauge, M.-T., Li, H., Racimo, F., Mallick, S., Schraiber, J. G., Jay, F., Prüfer, K., de Filippo, C., Sudmant, P. H., Alkan, C., Fu, Q., Do, R., Rohland, N., Tandon, A., Siebauer, M., Green, R. E., Bryc, K., … Pääbo, S. (2012b). A high-coverage genome sequence from an archaic Denisovan individual. *Science*, *338*(6104), 222–226.

Moreau, L. (2012a). Le Gravettien ancien d'Europe centrale revisité : mise au point et perspectives. In *L'Anthropologie* (Vol. 116, Issue 5, pp. 609–638). https://doi.org/10.1016/j.anthro.2011.10.002

Moreau, L. (2012b). Le Gravettien ancien d'Europe centrale revisité : mise au point et perspectives. *L'Anthropologie*, *116*(5), 609–638.

Morgan, C., Barton, L., Yi, M., Bettinger, R. L., Gao, X., & Peng, F. (2014). Redating Shuidonggou Locality 1 and Implications for the Initial Upper Paleolithic in East Asia. In *Radiocarbon* (Vol. 56, Issue 1, pp. 165–179). https://doi.org/10.2458/56.16270

Moroni, A., Ronchitelli, A., Arrighi, S., Aureli, D., Bailey, S., Boscato, P., Boschin, F., Capecchi, G., Crezzini, J., Douka, K., Marciani, G., Panetta, D., Ranaldo, F., Ricci, S., Scaramucci, S., Spagnolo, V., Benazzi, S., & Gambassini, P. (2018). Grotta del Cavallo (Apulia-Southern Italy). The Uluzzian in the mirror. *Journal of Anthropological Sciences = Rivista Di Antropologia: JASS / Istituto Italiano Di Antropologia*, *96*, 125–160.

Nejman, L., Wood, R., Wright, D., Lisá, L., Nerudová, Z., Neruda, P., Přichystal, A., & Svoboda, J. (2017). Hominid visitation of the Moravian Karst during the Middle-Upper Paleolithic transition: New results from Pod Hradem Cave (Czech Republic). *Journal of*



28

*Human Evolution*, *108*, 131–146.

Neruda, P., & Nerudová, Z. (2019). Technology of Early Szeletian leaf point shaping: a refitting approach. In *Archaeological and Anthropological Sciences* (Vol. 11, Issue 9, pp. 4515–4538). https://doi.org/10.1007/s12520-019-00818-3

Nerudová, Z., & Neruda, P. (2017). Technology of Moravian Early Szeletian leaf point shaping: A case study of refittings from Moravský Krumlov IV open-air site (Czech Republic). In *Quaternary International* (Vol. 428, pp. 91–108). https://doi.org/10.1016/j.quaint.2015.09.065

Pagani, L., Lawson, D. J., Jagoda, E., Mörseburg, A., Eriksson, A., Mitt, M., Clemente, F., Hudjashov, G., DeGiorgio, M., Saag, L., Wall, J. D., Cardona, A., Mägi, R., Wilson Sayres, M. A., Kaewert, S., Inchley, C., Scheib, C. L., Järve, M., Karmin, M., … Metspalu, M. (2016). Genomic analyses inform on migration events during the peopling of Eurasia. *Nature*, *538*(7624), 238–242.

Patterson, N., Moorjani, P., Luo, Y., Mallick, S., Rohland, N., Zhan, Y., Genschoreck, T., Webster, T., & Reich, D. (2012). Ancient admixture in human history. *Genetics*, *192*(3), 1065–1093.

Peng, F., Lin, S. C., Patania, I., Levchenko, V., Guo, J., Wang, H., & Gao, X. (2020). A chronological model for the Late Paleolithic at Shuidonggou Locality 2, North China. *PloS One*, *15*(5), e0232682.

Peresani, M., Bertola, S., Delpiano, D., Benazzi, S., & Romandini, M. (2019). The Uluzzian in the north of Italy: insights around the new evidence at Riparo Broion. In *Archaeological and Anthropological Sciences* (Vol. 11, Issue 7, pp. 3503–3536). https://doi.org/10.1007/s12520-018-0770-z

Pirson, S., Flas, D., Abrams, G., Bonjean, D., Court-Picon, M., Di Modica, K., Draily, C., Damblon, F., Haesaerts, P., Miller, R., Rougier, H., Toussaint, M., & Semal, P. (2012). Chronostratigraphic context of the Middle to Upper Palaeolithic transition: Recent data from Belgium. *Quaternary International: The Journal of the International Union for Quaternary Research*, *259*, 78–94.

Pitulko, V., Pavlova, E., & Nikolskiy, P. (2017). Revising the archaeological record of the Upper Pleistocene Arctic Siberia: Human dispersal and adaptations in MIS 3 and 2. *Quaternary Science Reviews*, *165*, 127–148.

Prüfer, K., Posth, C., Yu, H., Stoessel, A., Spyrou, M. A., Deviese, T., Mattonai, M., Ribechini, E., Higham, T., Velemínský, P., Brůžek, J., & Krause, J. (2021). A genome sequence from a modern human skull over 45,000 years old from Zlatý kůň in Czechia. *Nature Ecology & Evolution*. https://doi.org/10.1038/s41559-021-01443-x

Raghavan, M., Skoglund, P., Graf, K. E., Metspalu, M., Albrechtsen, A., Moltke, I., Rasmussen, S., Stafford, T. W., Jr, Orlando, L., Metspalu, E., Karmin, M., Tambets, K., Rootsi, S., Mägi, R., Campos, P. F., Balanovska, E., Balanovsky, O., Khusnutdinova, E., Litvinov, S., … Willerslev, E. (2014). Upper Palaeolithic Siberian genome reveals dual ancestry of Native Americans. *Nature*, *505*(7481), 87–91.

Reich, D., Patterson, N., Kircher, M., Delfin, F., Nandineni, M. R., Pugach, I., Ko, A. M.-S., Ko, Y.-C., Jinam, T. A., Phipps, M. E., Saitou, N., Wollstein, A., Kayser, M., Pääbo, S., & Stoneking, M. (2011). Denisova admixture and the first modern human dispersals into Southeast Asia and Oceania. *American Journal of Human Genetics*, *89*(4), 516–528.

Richter, D., Tostevin, G., & Skrdla, P. (2008). Bohunician technology and thermoluminescence dating of the type locality of Brno-Bohunice (Czech Republic). *Journal of Human Evolution*, *55*(5), 871–885.

Riel-Salvatore, J. (2009). What Is a “Transitional” Industry? The Uluzzian of Southern Italy as a Case Study. In *Sourcebook of Paleolithic Transitions* (pp. 377–396). https://doi.org/10.1007/978-0-387-76487-0_25

Riel-Salvatore, J., & Negrino, F. (2018). Proto-aurignacian lithic technology, mobility, and human niche construction: A case study from riparo bombrini, Italy. In *Studies in Human Ecology and Adaptation* (pp. 163–187). Springer International Publishing.

Roussel, M. (2013). Méthodes et rythmes du débitage laminaire au Châtelperronien : comparaison avec le Protoaurignacien. *Comptes rendus. Palevol*, *12*(4), 233–241.

Roussel, M., Soressi, M., & Hublin, J.-J. (2016). The Châtelperronian conundrum: Blade and


30

bladelet lithic technologies from Quinçay, France. *Journal of Human Evolution*, *95*, 13–32.

Ruebens, K., McPherron, S. J. P., & Hublin, J.-J. (2015). On the local Mousterian origin of the Châtelperronian: Integrating typo-technological, chronostratigraphic and contextual data. *Journal of Human Evolution*, *86*, 55–91.

Rybin, E. P. (2014). Tools, beads, and migrations: Specific cultural traits in the Initial Upper Paleolithic of Southern Siberia and Central Asia. *Quaternary International: The Journal of the International Union for Quaternary Research*, *347*, 39–52.

Sikora, M., Pitulko, V. V., Sousa, V. C., Allentoft, M. E., Vinner, L., Rasmussen, S., Margaryan, A., de Barros Damgaard, P., de la Fuente, C., Renaud, G., Yang, M. A., Fu, Q., Dupanloup, I., Giampoudakis, K., Nogués-Bravo, D., Rahbek, C., Kroonen, G., Peyrot, M., McColl, H., … Willerslev, E. (2019). The population history of northeastern Siberia since the Pleistocene. *Nature*, *570*(7760), 182–188.

Sinitsyn, A. A. (2007). Variabilité du Gravettien de Kostienki (Bassin moyen du Don) et des territoires associés. *Paléo*, *19*, 181–201.

Sinitsyn, A. A., & Hoffecker, J. F. (2006). Radiocarbon dating and chronology of the Early Upper Paleolithic at Kostenki. *Quaternary International: The Journal of the International Union for Quaternary Research*, *152-153*, 164–174.

Siska, V. (2019). *Human population history and its interplay with natural selection*. Apollo - University of Cambridge Repository. https://doi.org/10.17863/CAM.31536

Sitlivy, V., Chabai, V., Anghelinu, M., Uthmeier, T., Kels, H., Hilgers, A., Schmidt, C., Nita, L., Baltean, I., Veselsky, A., & Hauck, T. (2012). *The earliest Aurignacian in Romania: New investigations at the open air site of Româneşti-Dumbrăviţa I (Banat)*. https://doi.org/10.7485/QU59_4

Sitlivy, V., Chabai, V., Anghelinu, M., Uthmeier, T., Kels, H., Niţă, L., Băltean, I., Veselsky, A., & Ţuţu, C. (2014). Preliminary reassessment of the Aurignacian in Banat (South-western Romania). *Quaternary International: The Journal of the International Union for Quaternary Research*, *351*, 193–212.



31

Skoglund, P., Mallick, S., Bortolini, M. C., Chennagiri, N., Hünemeier, T., Petzl-Erler, M. L., Salzano, F. M., Patterson, N., & Reich, D. (2015). Genetic evidence for two founding populations of the Americas. *Nature*, *525*(7567), 104–108.

Škrdla, P. (2017). Middle to Upper Paleolithic transition in Moravia: New sites, new dates, new ideas. In *Quaternary International* (Vol. 450, pp. 116–125). https://doi.org/10.1016/j.quaint.2016.07.029

Slavinsky, V. S., Rybin, E. P., Khatsenovich, A. M., & Belousova, N. E. (2019). Intentional fragmentation of blades in the initial upper Paleolithic industries of the Kara-Bom site (Altai, Russia). In *Archaeological Research in Asia* (Vol. 17, pp. 50–61). https://doi.org/10.1016/j.ara.2018.05.002

Tafelmaier, Y. (2017). *Technological Variability at the Beginning of the Aurignacian in Northern Spain: Implications for the Proto- and Early Aurignacian Distinction*.

Teyssandier, N., Bon, F., & Bordes, J.-G. (2010a). WITHIN PROJECTILE RANGE: Some Thoughts on the Appearance of the Aurignacian in Europe. In *Journal of Anthropological Research* (Vol. 66, Issue 2, pp. 209–229). https://doi.org/10.3998/jar.0521004.0066.203

Teyssandier, N., Bon, F., & Bordes, J.-G. (2010b). WITHIN PROJECTILE RANGE: Some Thoughts on the Appearance of the Aurignacian in Europe. In *Journal of Anthropological Research* (Vol. 66, Issue 2, pp. 209–229). https://doi.org/10.3998/jar.0521004.0066.203

Tostevin, G. B. (2003). Attribute analysis of the lithic technologies of stránská skála ii-iii in their regional and inter-regional context: Origins of the upper paleolithic in the brno basin. In *Stránská skála: Origins of the upper paleolithic in the brno basin* (pp. 77–118). Peabody Museum Press, Harvard University.

Trinkaus, E., & Buzhilova, A. P. (2018). Diversity and differential disposal of the dead at Sunghir. *Antiquity*, *92*(361), 7–21.

Trinkaus, E., Buzhilova, A. P., Mednikova, M. B., & Dobrovol'skai͡a, M. V. (2014). *The People of Sunghir: Burials, Bodies, and Behavior in the Earlier Upper Paleolithic*. Oxford University Press.

Tsanova, T. (2008). *Les débuts du Paléolithique supérieur dans l'Est des Balkans: Réflexion*

 32

*à partir de l'étude taphonomique et techno-économique des ensembles lithiques des sites de Bacho Kiro (couche 11), Temnata (couches VI et 4) et Kozarnika (niveau VII)*. https://doi.org/10.30861/9781407301914

Usik, V. I., Monigal, K., & Kulakovskaya, L. (2006). *New perspectives on the Transcarpathian Middle to Upper Paleolithic boundary*. na.

Vishnyatsky, L. B., & Nehoroshev, P. E. (2004). The beginning of the Upper Paleolithic on the Russian Plain. *The Early Upper Paleolithic Beyond Western Europe*, 80–96.

Wall, J. D. (2017). Inferring Human Demographic Histories of Non-African Populations from Patterns of Allele Sharing. *American Journal of Human Genetics*, *100*(5), 766–772.

Yang, M. A., Gao, X., Theunert, C., Tong, H., Aximu-Petri, A., Nickel, B., Slatkin, M., Meyer, M., Pääbo, S., Kelso, J., & Fu, Q. (2017). 40,000-Year-Old Individual from Asia Provides Insight into Early Population Structure in Eurasia. *Current Biology: CB*, *27*(20), 3202–3208.e9.

Zilhão, J., d'Errico, F., Bordes, J.-G., Lenoble, A., Texier, J.-P., & Rigaud, J.-P. (2006). Analysis of Aurignacian interstratification at the Chatelperronian-type site and implications for the behavioral modernity of Neandertals. *Proceedings of the National Academy of Sciences of the United States of America*, *103*(33), 12643–12648.

Zwyns, N., & Lbova, L. V. (2019). The Initial Upper Paleolithic of Kamenka site, Zabaikal region (Siberia): A closer look at the blade technology. *Archaeological Research in Asia*, *17*, 24–49.

Zwyns, N., Paine, C. H., Tsedendorj, B., Talamo, S., Fitzsimmons, K. E., Gantumur, A., Guunii, L., Davakhuu, O., Flas, D., Dogandžić, T., Doerschner, N., Welker, F., Gillam, J. C., Noyer, J. B., Bakhtiary, R. S., Allshouse, A. F., Smith, K. N., Khatsenovich, A. M., Rybin, E. P., … Hublin, J.-J. (2019). The Northern Route for Human dispersal in Central and Northeast Asia: New evidence from the site of Tolbor-16, Mongolia. *Scientific Reports*, *9*(1), 11759.

Zwyns, N., Rybin, E. P., Hublin, J.-J., & Derevianko, A. P. (2012). Burin-core technology and laminar reduction sequences in the initial Upper Paleolithic from Kara-Bom (Gorny-Altai,



33

Siberia). In *Quaternary International* (Vol. 259, pp. 33–47). https://doi.org/10.1016/j.quaint.2011.03.036

 34
